# The transcription factor Traffic jam orchestrates the somatic piRNA pathway in *Drosophila* ovaries

**DOI:** 10.1101/2024.09.10.612307

**Authors:** Azad Alizada, Aline Martins, Nolwenn Mouniée, Julia V. Rodriguez Suarez, Benjamin Bertin, Nathalie Gueguen, Vincent Mirouse, Anna-Maria Papameletiou, Austin J. Rivera, Nelson C. Lau, Abdou Akkouche, Stéphanie Maupetit-Mehouas, Gregory J. Hannon, Benjamin Czech Nicholson, Emilie Brasset

**Author notes:** These authors contributed equally to this work (Azad Alizada, Aline Martins, Nolwenn Mouniée, Julia V. Rodriguez Suarez). Corresponding authors: GJH, BCN, EB.

## Abstract

The PIWI-interacting RNA (piRNA) pathway is essential for transposable element (TE) silencing in animal gonads. While the transcriptional regulation of piRNA pathway components in germ cells has been documented in mice and flies, their control in somatic cells of *Drosophila* ovaries remains unresolved. Here, we demonstrate that Traffic jam (Tj), the *Drosophila* orthologue of large Maf transcription factors in mammals, is a master regulator of the somatic piRNA pathway. Tj binds to regulatory regions of somatic piRNA factors and the major piRNA cluster *flamenco*, which carries a Tj-bound enhancer downstream of its promoter. Depletion of Tj in somatic follicle cells causes downregulation of piRNA factors, loss of *flam* expression and de-repression of *gypsy*-family TEs. We propose that the arms race between the host and TEs led to the co-evolution of promoters in piRNA pathway genes as well as TE regulatory regions that both rely on a shared transcription factor.

**Highlights:**

- Traffic jam (Tj) acts as a master regulator of the somatic piRNA pathway in *Drosophila*.
- Tj regulates a network of piRNA pathway genes, mirroring the gene-regulatory mechanism of A-MYB in the mouse testis and Ovo in fly ovaries.
- *Cis*-regulatory elements with Tj motifs are present at the promoters of somatic piRNA pathway genes.
- The expression of the *flamenco* piRNA cluster is directly controlled by Tj.

## Introduction

Transposable elements are DNA sequences with the ability to move or replicate to new positions within the host genome^1^. Succeeding waves of TE mobilization and repression have allowed TEs to accumulate in the genomes of nearly all organisms. Approximately 20% of the *Drosophila* genome and more than 50% of the human genome is composed of TE sequences^2^. While their ability to transpose makes them powerful facilitators of genome evolution, their mobilization can also have deleterious effects, such as mutations, gene disruptions, and chromosomal rearrangements^3,4^. The resulting selective pressure has driven the co-evolution of many mechanisms within the hosts to effectively counter the disruptive TE activity^5^.

In animals, the piRNA pathway is one of the key defence mechanisms against active TEs. In *Drosophila*, the piRNA pathway is crucial for maintaining genome stability, particularly in gonadal tissues, which consist of germ cells surrounded by somatic follicle cells^6–8^. Previous work has uncovered the existence of distinct piRNA pathways in germ cells and somatic follicle cells^7–9^. Germ cells express a more sophisticated piRNA system that involves both post-transcriptional silencing of transposon transcripts via the ping-pong loop and co-transcriptional silencing of nascent TEs. The first mechanism requires Aubergine, Argonaute-3 and numerous Tudor and RNA binding proteins such as Qin, Tejas, Vasa, Krimper, Spn-E and BoYb, whereas co-transcriptional TE silencing relies on Piwi, its cofactor Arx and the PICTS complex composed of Panx, Nxf2, Nxt1, and Ctp^10–16^. Furthermore, piRNAs in germ cells are predominantly generated from precursor transcripts derived from discrete genomic loci called dual-strand piRNA clusters^6,17–19^. The target repertoires of these piRNA clusters are dictated by both full-length and truncated transposon copies and their transcription and export relies on non-canonical mechanisms, involving Rhino (Rhi), Deadlock (Del), Cutoff (Cuff), Moonshiner (Moon), Nxf3, and Bootlegger (Boot)^6^.

To avoid the robust silencing machinery in germ cells, some TEs such as *gypsy* or ZAM have gained the ability to express exclusively in the somatic follicle cells, form virus-like particles and invade germ cells such as the oocyte^20–22^. This infectious capacity enables their transposition into the germline genome and hence transmission of new TE copies to future generations. As a consequence of this host-transposon arms race, somatic follicle cells have adapted a simplified piRNA pathway version comprised of the factors required for piRNA biogenesis (e.g., Armi, Zuc, Mino, Gasz, Daed, SoYb, Vret, Shu and the soma-specific Yb) and co-transcriptional transposon silencing (e.g., Piwi, Arx, Panx, Nxf2, Nxt1, and Ctp)^8,13,14,23–27^. In addition, somatic piRNA clusters have emerged to control TE expression in the somatic follicle cells of the ovary. In contrast to the germline-active dual-strand piRNA clusters, the soma-expressed piRNA clusters are unistrand and give rise to piRNAs capable of targetting active TEs. The major somatic piRNA cluster in *Drosophila* ovaries is *flamenco* (*flam*)^6,28^. Contrary to the dual-strand piRNA clusters, *flam* is conserved across millions of years of Drosophilid evolution^29^, and transcribed by a canonical mechanism that includes a promoter, splicing, and polyadenylation^30^. Furthermore, to-date, *flam* is the only known essential piRNA cluster, with *flam* mutants showing rudimentary ovaries and complete female sterility^18,31^.

Despite the importance of the somatic piRNA pathway, including the *flam* cluster and the somatic piRNA factors, the gene-regulatory circuit controlling their transcription remains to be determined. A recent study has shown that the transcription factor (TF) Ovo plays a key role in coordinating the expression of germline-specific piRNA pathway genes in the ovaries^32^. However, it remains unclear whether one or multiple TF(s) drive the somatic piRNA pathway in the ovarian follicle cells. Here, we uncover Traffic jam (Tj), a large Maf TF, as a master regulator of the somatic piRNA pathway. Tj is critical for gonad development^33^ and was previously implicated in the regulation of *piwi* expression in somatic cells^34^. In this study, we extend these findings by showing that Tj regulates the expression of many somatic piRNA pathway factors, as well as the key unistrand piRNA cluster, *flamenco*. We demonstrate that Tj directly binds to the regulatory regions of these genes, driving their soma-specific expression in ovarian follicle cells. Furthermore, we identify a specific enhancer region downstream of the *flam* TSS that is essential for its transcriptional activity in somatic cells and harbors Tj binding sites. Our findings shed light on the intricate transcriptional control necessary for TE silencing in the ovarian soma, offering new insights into the regulation of the soma-specific piRNA pathway.

## Results

### The transcription factor Traffic jam controls somatic piRNA factor expression

To uncover the regulatory mechanisms underlying the expression of somatic piRNA pathway factors, we first performed differential gene expression analysis between germline and somatic cells derived from *Drosophila* ovaries. We dissected and dissociated ovaries of transgenic *D. melanogaster* flies expressing GFP specifically in germ cells (using the *vasa* promoter) and sorted the cells using FACS into germline (*vas*-GFP+) and somatic (*vas*-GFP-) populations^32^. We then extracted RNAs and generated RNA-seq libraries before carrying out a differential gene expression analysis between germline and somatic cells. This approach identified piRNA pathway genes with significant expression in the soma of the *Drosophila* ovary, namely*, fs(1)Yb*, *nxf2*, *panx*, and *armi* (**Figure 1A, left**). Among these factors, *fs(1)Yb* was reported to be exclusively expressed in somatic cells^35^, while the other factors are known to also function in germ cells^36^. As expected these four soma-enriched piRNA factors also showed expression in ovarian somatic cells (OSCs) (**Figure 1A**), a cell line derived from ovary somatic follicle cells^34,37^.

**Figure 1:**
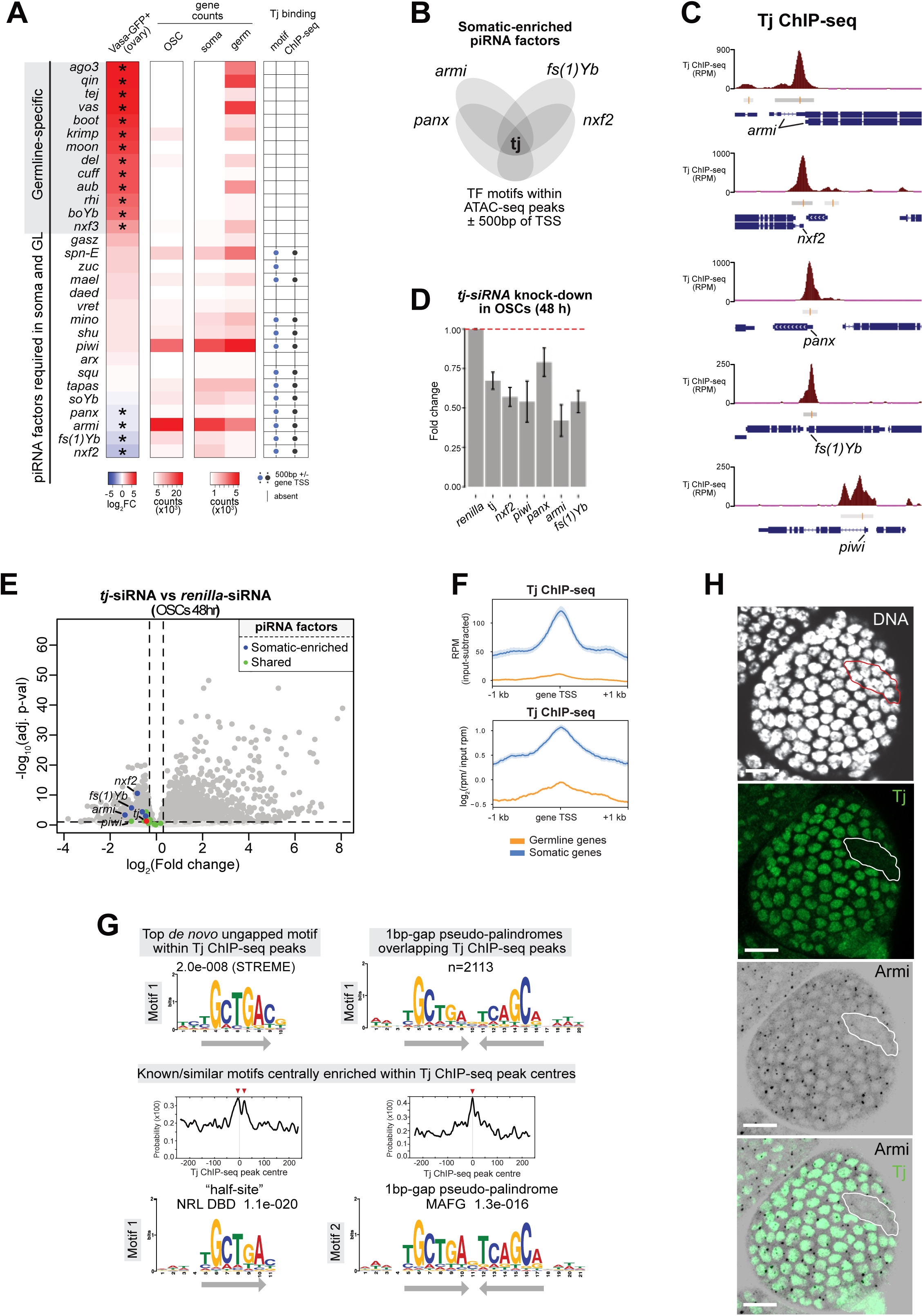
Traffic jam regulates the expression of somatic piRNA pathway genes in *Drosophila* ovarian somatic cells. **(A)** Heatmap showing germline-specific and soma-expressed piRNA pathway genes ranked by gene expression fold-enrichments in the FACS-sorted germline cells (*vas*-GFP+) compared to somatic cells (*vas*-GFP-) dissociated from fly ovaries (*<0.01, DEseq2, RNA-seq n=3 replicates from distinct samples). Normalized RNA-seq counts (DEseq2) from OSCs, vas-GFP+ germline cells, and vas-GFP-somatic cells are also shown. The presence of Tj motifs within ovary ATAC-seq peaks and Tj ChIP-seq peaks (ENCODE, female whole-fly) within ±500 bp of gene TSS is depicted with dots. **(B)** Venn diagram showing Tj as the only motif shared between OSC ATAC-seq peaks of the promoter regions (±500 bp of TSS) of somatic-enriched piRNA pathway genes. Full TF motif list is shown in **Table S1**. **(C)** Tj ChIP-seq from whole female flies showing Tj binding events to the promoters of somatic piRNA pathway genes (dm6). Grey bars indicate Tj ChIP-seq peaks with the central red lines showing peak summits. **(D)** RT-qPCR plot showing 48 h siRNA knockdown of *tj* in OSCs resulting in downregulation of somatic piRNA pathway genes (normalised to *rpL32*; n=3 replicates from distinct samples; error bars indicate standard deviation). **(E)** Volcano plot showing differential RNA-seq analysis (DEseq2) between *tj* and *renilla* siRNA knockdowns (48 h; n=3 replicates from distinct samples). Blue dots are showing soma-enriched piRNA pathway genes (*fs(1)Yb*, *nxf2*, *panx*, *soYb* and *armi*); green dots showing general piRNA factors (e.g., *piwi*); red dots are showing *tj*. **(F)** Genomic binding profiles of Tj at the promoters of germline- and soma-enriched genes using normalized ChIP-seq (ENCODE; whole-fly; n=3) signals with input subtraction (top) and log2 of chip rpm/input rpm (bottom). **(G)** The top-scoring *de novo* motif within Tj ChIP-seq peaks identified using STREME (MEME-ChIP) (top-left). The numbers and motif logo (MEME) of genomic sites matching the 1-bp gapped pseudo-palindromic motif (MA0659.1; JASPAR 2024) within Tj ChIP-seq peaks identified using FIMO tool (p<1x10^-04^) (top-right). Enrichment of the known vertebrate motifs within Tj ChIP-seq peak centers using CentriMo (MEME-ChIP) (bottom). **(H)** Confocal images of *tj^eo^*^2^ clonal egg chamber showing Armi (black) and Tj (green) protein expression via immunostaining. Clone is circled in white. Nuclei are labelled with DAPI (white). Scale bars: 10 µm.

Next, to identify the potential TFs controlling the expression of these somatic-enriched piRNA factors (*fs(1)Yb, nxf2, panx*, and *armi*), we analysed ATAC-seq from wildtype OSCs^38^ and performed motif scanning using the FIMO tool (match p<0.001; in MEME suite) to look for transcription factor motifs (the FlyFactorSurvey database) that were shared between their promoter ATAC-seq peaks (±500 bp of the transcription start sites) (**Figure 1B and Table S1**). Surprisingly, this analysis identified only a single transcription factor motif shared between the promoter ATAC-seq peaks of the four soma-enriched piRNA factors, Tj. We confirmed *in vivo* Tj binding to the promoters of *fs(1)Yb*, *nxf2*, *panx*, and *armi* (**Figure 1C**) by re-analysing publicly available Tj-GFP ChIP-seq (referred to as Tj ChIP-seq) data from whole female flies^39^. Moreover, Tj ChIP-seq peaks and Tj motifs were also present within promoters of other somatic piRNA factors that are expressed in both somatic and germline cells such as *piwi* but these were absent in promoters of the germline-specific piRNA factors (**Figure 1A, right and Figure 1C**), suggesting that Tj could be specifically controlling somatic piRNA pathway genes.

To validate our hypothesis that Tj regulates soma-expressed piRNA factors, we performed *tj* knockdowns in OSCs, followed by RT-qPCR. Tj depletion resulted in downregulation of somatic piRNA factors (**Figure 1D**). Although *tj* levels were reduced by ∼40% following 48 hours of knockdown, we observed similar levels of downregulation for *nxf2*, *piwi*, *armi*, and *fs(1)yb* (RT-qPCR, n=3 replicates from distinct samples, p<0.01) (**Figure 1D**). To corroborate these results, we performed RNA-seq following 48 hours and 96 hours of *tj* knockdown. These also showed significant downregulation of the soma-enriched piRNA factors *fs(1)Yb*, *nxf2*, *panx*, *soYb*, and *armi* (RNA-seq, n≥3 replicates from distinct samples, p.adj.<0.01) (**Figure 1E and Figure S1A**). The piRNA factors shared between the somatic and germline cells which showed Tj binding at their promoters *in vivo* (i.e., *piwi*, *vret*, *mino*, and *shu*), were similarly downregulated upon *tj* knockdowns in OSCs (**Figures 1E and S1A**). These results suggest that Tj is a soma-enriched TF responsible for the regulation of soma-expressed piRNA pathway genes in *Drosophila* ovarian somatic cells, likely via direct promoter binding.

Genome-wide Tj ChIP-seq analysis revealed 7,956 Tj binding sites overall, which were enriched at the promoters of soma-expressed genes (including the somatic piRNA pathway genes), and depleted from promoters of the germline-enriched genes, including germline-specific piRNA pathway genes (**Figures 1A and 1F**). Specifically, Tj peaks overlapped with 714 transcription start sites (within ±500 bp of TSS) of 419 soma-expressed genes (defined using RNA-seq log2FC<-1, adjusted p.value<0.01, n=3 from FACS-sorted *vas*-GFP-cells), representing 62% of all soma-enriched gene TSSs (n=1,142) (**Table S1**). Motif analysis revealed that the top-scoring *de novo* motif within Tj ChIP-seq peaks had a consensus sequence of TGCTGAC (STREME e-value=2.0x10^-08^). We noticed that the top-enriched motif within the peak centers had a consensus sequence of TGCTGA (CentriMo e-value=1.1x10^-20^) and was often found nearby its reverse complement TCAGCA (as close as by 1 bp spacing), resulting in a pseudo-palindromic TGCTGA(N) nTCAGCA motif (CentriMo e-value=1.3x10^-16^; FIMO n=2,113 sites) (**Figures 1G and S1B**). Interestingly, this closely mirrors the Maf recognition element (MARE) motifs recognized by DNA-binding domains of the dimeric basic Leucine Zipper Maf family transcription factors^40,41^, suggesting that Tj could bind some target sites as a dimer. Overall, our results indicate that Tj specifically directs transcriptional regulation of the somatic ovarian program, including somatic piRNA pathway components, by binding to TF motifs at the promoters of somatic target genes.

To test Tj’s role in regulating somatic piRNA factors in an *in vivo* setting, we disrupted Tj activity and examined piRNA factors expression through immunostaining in the *Drosophila* adult ovaries (**Figure 1H**). As Tj plays a critical role in regulating the formation and function of the germline stem cell niche and its mutation leads to ovarian atrophy^42,43^, we abolished Tj activity in mosaic clones of follicle cells carrying a genetic null allele of Tj, the *tj^eo^*^2^ allele^33^. This allele carries a point mutation resulting in a premature stop codon resulting in a truncated version of the Tj protein, which lacks critical DNA-binding domains, including the leucine zipper and nuclear localization signal^33^. As expected, in wild-type follicle cells, Armi, protein essential for piRNA biogenesis, accumulate in distinct foci representing Yb-bodies near the nuclear periphery^44^. However, in *tj* mutant clones, Armi proteins were completely absent (**Figure 1H**). This finding confirms that, as observed in OSCs, Tj regulates piRNA factor expression within follicle cells of the adult ovary.

### Tj binds to conserved motifs in the regulatory region of the piRNA cluster *flam*

We next asked whether Tj also regulates somatic piRNA cluster transcription. The major piRNA cluster active in OSCs and somatic follicle cells is *flamenco*^6,28^. Analysis of ovary and OSC ATAC-seq data^29,38^ revealed peaks upstream of *flam* that correspond to the nearby gene DIP1 (ATAC-seq peaks #1-4) as well as three conserved peaks 1 kb upstream and 5 kb downstream of the *flam* TSS in *D. melanogaster* (ATAC-seq peaks #5, #6 and #7) that were also present in the orthologous region in the *D. yakuba* genome (**Figure 2A**). Peak #5 harbored the motif of cubitus interruptus, a TF previously reported to contriunte to *flam* expression^45^, while peak #6 contained the initiator element (Inr) of the *flam* TSS^45^. Peak #8 overlapped an alternative, further downstream TSS that was previously identified by PRO-seq (the second *flam* PRO-seq transcription initiation peak^46^. We noted that the conserved peaks #5 and #7 had disproportionately stronger ATAC-seq signals in OSCs compared to the whole ovaries, suggesting their somatic enrichment (**Figure 2A**). To identify enhancer activity genome-wide, we re-analysed Self-Transcribing Active Regulatory Region sequencing (STARR-seq) data performed in OSCs using genomic DNA of both *D. melanogaster* and *D. yakuba*^47–49^. We found that enhancer activity signal overlapped with ATAC-seq peak #7 in *D. melanogaster* and its corresponding peak in the *D. yakuba* (**Figure 2A**). Motif scanning (FIMO) revealed 21 transcription factor motifs (the FlyFactorSurvey database) that were shared between peak #7 in *D. melanogaster* and its orthologous region in *D. yakuba* **(Figure 2B)**. Intriguingly, the motif for Tj was among the shared hits. Expression filtering using RNA-seq data from ovaries of *D. melanogaster* and *D.yakuba* revealed TF candidates expressed in this tissue in both species **(Figure 2B)**. Of these, four (*tj*, *crc*, *br*, and *vri*) showed soma-enrichment with *tj* being the top hit **(Figure 2C)**.

**Figure 2:**
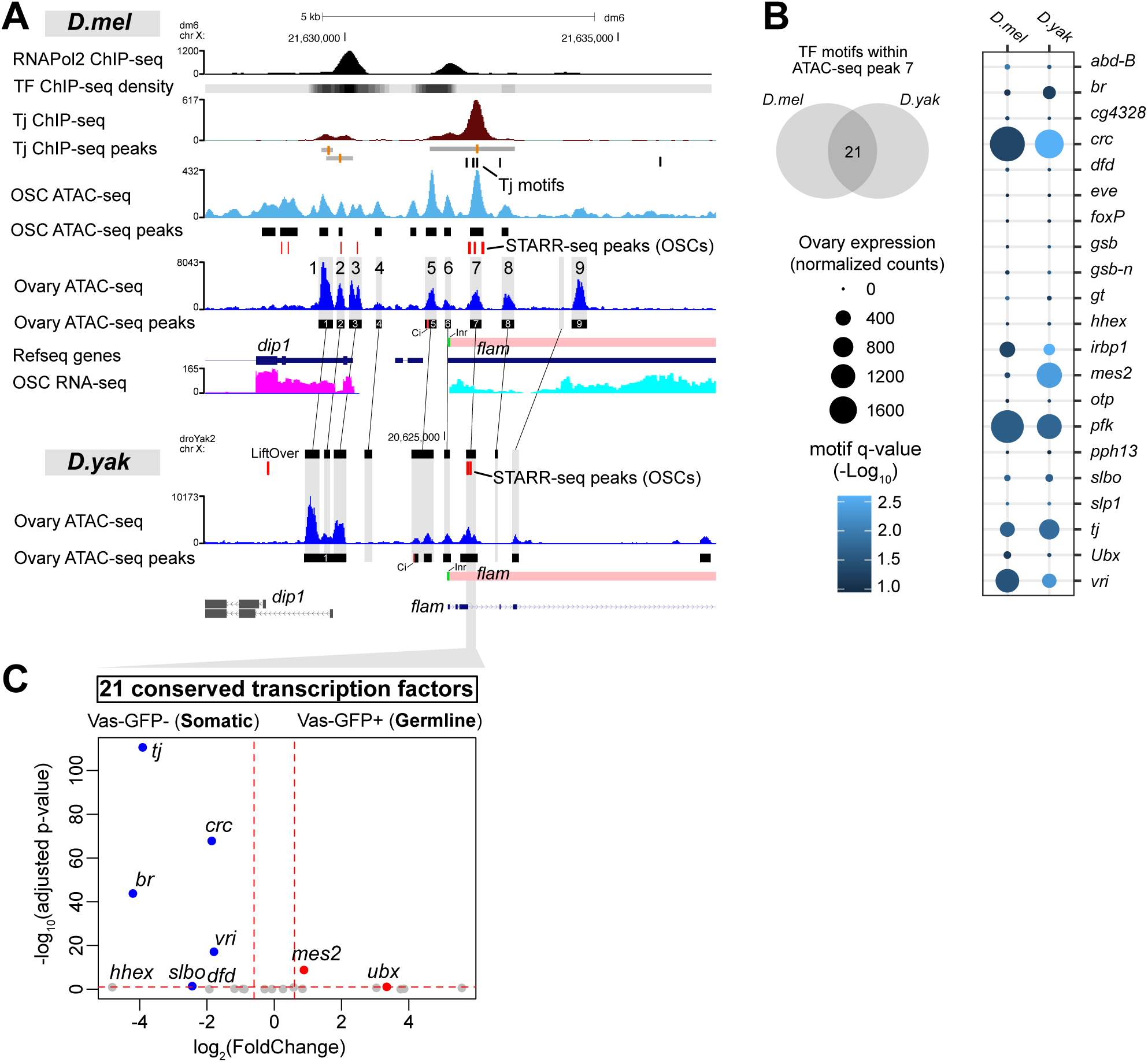
Tj binds to conserved *cis*-regulatory elements in *flamenco*. **(A)** Conservation of ovary ATAC-seq peaks in the *flam* promoter region of *D. melanogaster* (*dm6*) and *D. yakuba* (*droYak2*) species (orthologous regions derived with UCSC LiftOver and chain files). OSC ATAC-seq peaks, OSC STARR-seq peaks, OSC RNA-seq, RNAPol2 ChIP-seq (ovary)^66^ and Tj ChIP-seq (ENCODE, whole-fly) are shown. **(B)** Venn diagram showing motifs for 21 TFs shared between ATAC-seq peak #7 in *D. melanogaster* and its orthologous peak region in *D. yakuba*. Dot plot showing expression of 21 candidate TFs in the ovaries of *D. melanogaster* and *D. yakuba* (read depth and gene length-normalized DEseq2 counts, RNA-seq n=4). **(C)** Soma and germline-enrichment of 21 candidate TFs based on differential gene expression between somatic (*vas*-GFP-) and germline (*vas*-GFP+) cells from the FACS-sorted fly ovaries (DEseq2 was performed between *vas*-GFP-(n=3) and *vas*-GFP+ (n=3) RNA-seq libraries using counts from all genes).

In the *D. melanogaster* genome, Tj motifs were found downstream of the *flam* TSS at positions +327 to +332 bp (chrX:21,632,218-21,632,223; dm6), +437 to +450 bp (mutated pseudo-palindrome; chrX:21,632,328-21,632,340; dm6), +517 to +523 bp (summit of Tj ChIP-seq peak; chrX:21,632,408-21,632,414; dm6), and +935 to +940 bp (upstream of the alternative *flam* TSS; chrX: 21,632,826-21,632,831; dm6) (**Figure 2A**). Through Tj ChIP-seq analysis, we confirmed a strong Tj binding peak (-336 bp to +1214 bp) ovarlapping these motifs. Interestingly, two of the Tj motifs (+437 to +450 bp and +517 to +523 bp) also overlapped the conserved ovary ATAC-seq peak #7 (**Figure 2A**). The summits of the Tj ChIP-seq peak, the ovary ATAC-seq peak and the OSC ATAC-seq peak coincided with the +517 to +523 bp Tj motif (**Figure 2A**). Taken together, our results suggest that, in addition to regulation of somatic piRNA genes, Tj may also control the piRNA cluster *flam*.

### Expression of *flam* in follicle cells is dependent on a regulatory region located downstream of its TSS

Previous work identified the major TSS, and the minimal promoter of *flam* in OSS cells^45^. To confirm the *flam* regulatory regions, including putative Tj motifs, *in vivo*, we generated transgenic flies carrying transgene expressing the *tomato* reporter gene under the control of different *flam* promoter regions (starting at -1626 bp upstream and ending at +6334 bp downstream of the *flam* TSS) (**Figure 3A**). To compare the relative expression of these reporter constructs, we integrated all reporter variants into the identical genomic site at cytological position 53B2 and followed Tomato expression in fly ovaries by immunostaining and RT-qPCR (**Figures 3A-C**). The strongest Tomato expression in somatic follicle cells was observed for the -515; +1200 and the -515; +2086 constructs, with weaker signals found with the -515; +718 construct (**Figures 3B-C**). The increased Tomato expression observed in the -515; +1200 line prompted us to further explore this region. The presence of the fourth Tj motif identified at +935 to +940 bp (TCAGCA) together with the alternative *flam* TSS immediately downstream of this region at +977 to +1202 (ovary ATAC-seq peak #8 matching the second *flam* PRO-seq transcription initiation peak) potentially contributed to the enhanced Tomato expression observed when the region in constructs extending beyond +718.

**Figure 3:**
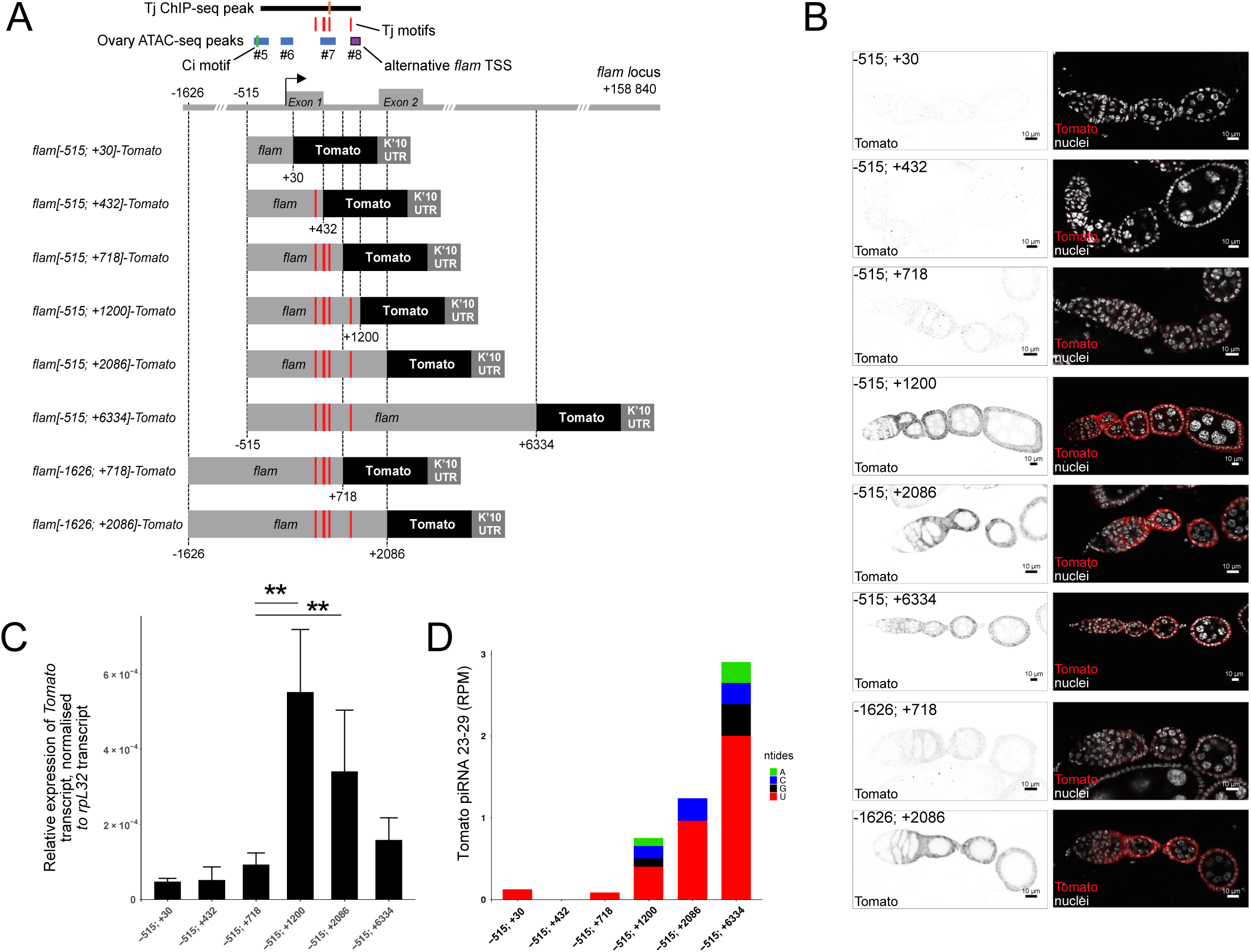
An enhancer of *flam* is located downstream of the TSS. **(A)** Schematic representation of the transgenic Tomato reporter constructs. The positions of the Tj ChIP-seq peak (line in the middle showing the peak summit), Tj motifs (red) and ATAC-seq peaks (#5-8) are highlighted above. **(B)** Confocal images of ovarioles of the indicated reporters showing on the left panels Tomato immunostaining (grey) and on the right panels the merge of Tomato immunostaining (red) and nuclei stained with DAPI (white). Scale bars: 10 µm. **(C)** Relative transgene expression levels in ovaries measured by RT-qPCR. *tomato* expression was normalized to *rpl32*. Mean expression is shwon (n=6) with error bars representing standard deviation. Statistical significance was determined using the Mann-Whitney test (** indicating a P-value < 0.005 and * a P-value < 0.05). **(D)** Amount of piRNAs produced from the *tomato* reporter sequence, expressed in reads per million (RPM). Nucleotide bias of the first nucleotide of piRNAs are shown as indicated.

Surprisingly almost no artificial tomato piRNAs was detected with the -515; +432 and - 515; +718 constructs (**Figure 3D**). These results suggest that the first 718 bp of the *flam* transcript, which were previously described as the piRNA trigger sequence in OSS cells^50^, are not sufficient to direct artificial piRNA processing from the *tomato* reporter *in vivo* (**Figure 3D**). In contrast, a longer -515; +6334 construct showed a weaker Tomato expression but exhibited a markedly stronger piRNA production relative to the -515; +1200 and -515; +2086 reporters (**Figures 3B-D**). Hence, including more *flam* sequence appeared to enhance piRNA processing efficiency and resulted in a concordant decrease of reporter mRNAs. This observation could be similar to the size-dependent dynamics observed in pachytene piRNA clusters in mice^51^. As expected, no Tomato expression was detected in the germline regardless of the constructs used (**Figure 3B**).

Neither the transgene carrying the -515; +30 *flam* regulatory region (which includes the minimal promoter), nor the reporter comprising -515; +432 (which includes the first exon) was sufficient to drive strong Tomato expression in somatic follicle cells (**Figure 3B-C**). These results suggest that while the Ci binding site (-394 to -366 bp from the *flam* TSS) is required for minimal *flam* expression in OSS cells, it is unable to induce strong expression in adult follicle cells. To confirm that the minimal *flam* promoter we have previously identified can efficiently drive transcription *in vivo,* we added UAS sequences in front of the -515; +432 *flam* promoter (**Figure 4A**). When combined with a follicle cell-specific *traffic jam* (*tj*)-Gal4 driver, we indeed observed strong Tomato expression in somatic follicle cells, (**Figures 4B-C**). Next, using CRISPR/Cas9, we deleted the minimal promoter region and tested its requirement for *flam* expression (**Figures 4A,D-F**). The resulting *flam* mutants, named *flamΔprom*, were female sterile and exhibited fully atrophied ovaries, along with a loss of *flam* piRNAs, while the expression of the germline piRNA clusters remained unaffected (**Figure 4D-F**). These results suggest that the -515 and +432 region (relative to the *flam* TSS) contains the minimal promoter necessary for *flam* expression but not its full regulatory region. Furthermore, no Tomato expression was detected in ovaries of transgenic flies carrying the +770; +2086 *flam* region (**Figures S2A and S2B**). Overall, our findings suggest that the sequence between +432 and +1200 bp downstream of the *flam* TSS that overlaps Tj ChIP-seq peaks and contains Tj motifs, functions as an enhancer driving efficient soma-specific *flam* expression in the adult ovary.

**Figure 4:**
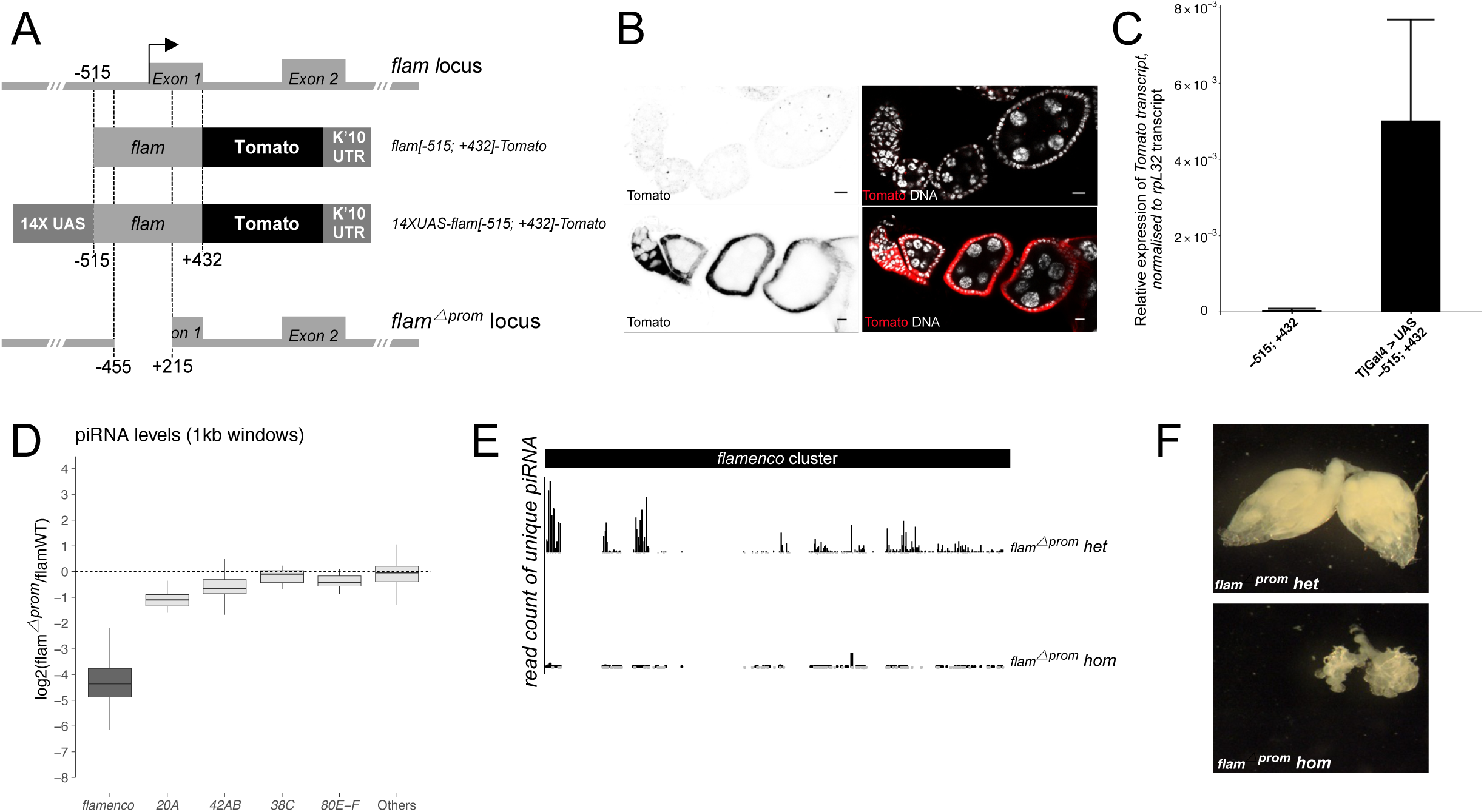
The minimal *flam* promoter is crucial for its expression and piRNA biogenesis. **(A)** Schematic representation of the reporter constructs and *flam* promoter deletion (*flam*△*prom* locus). **(B)** Confocal images of ovarioles of the indicated reporters showing Tomato (red) expression by immunostaining. Nuclei are labelled with DAPI (white). Scale bars: 10 µm. **(C)** Relative transgene expression levels in ovaries measured by RT-qPCR. Tomato expression was normalized to *rpL32*. Mean expression is shwon (n=6), with error bars representing standard deviation. Statistical analysis was performed using the Mann-Whitney test (** indicates P-value of 0.005). **(D)** Boxplot showing changes in piRNA levels for the indicated clusters (split into 1kb tiles) in *flam* mutants (*flam*△*prom*) relative to controls (*flam* WT). The boxplot displays the median (line) and the first and third quartiles (box). **(E)** Genome uniquely mapping piRNAs (normalized per million of genome-mapped reads with 0 mismatches) are shown from heterozygous (top) and homozygous *flam* mutant (bottom) ovaries. **(F)** Morphology of *Drosophila* ovaries from *flam* heterozygous (top) and homozygous mutant (bottom) flies.

We also investigated whether the sequence upstream of the *flam* TSS influences its expression by testing a larger *flam* promoter that includes the -1624 bp region that ranges from the *flam* TSS to the closest upstream gene, *DIP1* (**Figure 3A**). This extended promoter did not enhance Tomato expression in the somatic follicle cells of the -1624; +718 and -1624; +2086 transgenic lines (**Figures 3A-C**). These findings suggest the absence of additional enhancers upstream of the minimal *flam* promoter.

### Tj is the master transcription factor for follicle cell-specific expression of *flam*

To test for a direct regulation of the *flam* piRNA cluster by Tj *in vivo*, we abolished Tj activity and monitored *flam* expression by smFISH in mosaic clones of follicle cells (**Figures 5A and S3**). We observed a complete absence of *flam* signal in *tj*-deficient mutant cells, whereas *flam* signal at the nuclear periphery was detected in neighboring control cells that were Tj positive, consistent with previous reports^52,53^. Furthermore, sequencing of small RNAs from ovaries with soma-specific knockdown (SKD) of Tj (using a *tj*-Gal4 driver), showed a complete loss of piRNAs from the *flam* locus. (**Figure 5B**). Since Tj has been previously reported to regulate Piwi ^34^ and to rule out indirect effects, we investigated *flam* transcription by smFISH in follicle cells upon SKD of *piwi*. In contrast to *tj* mutant follicle cells, which lack *flam* expression, the absence of Piwi had no effect on *flam* levels (**Figures 5C-D**). In conclusion, our findings strongly support a key role of Tj in directly regulating the transcription of *flam* piRNA precursors in the somatic follicle cells of the adult ovary.

**Figure 5:**
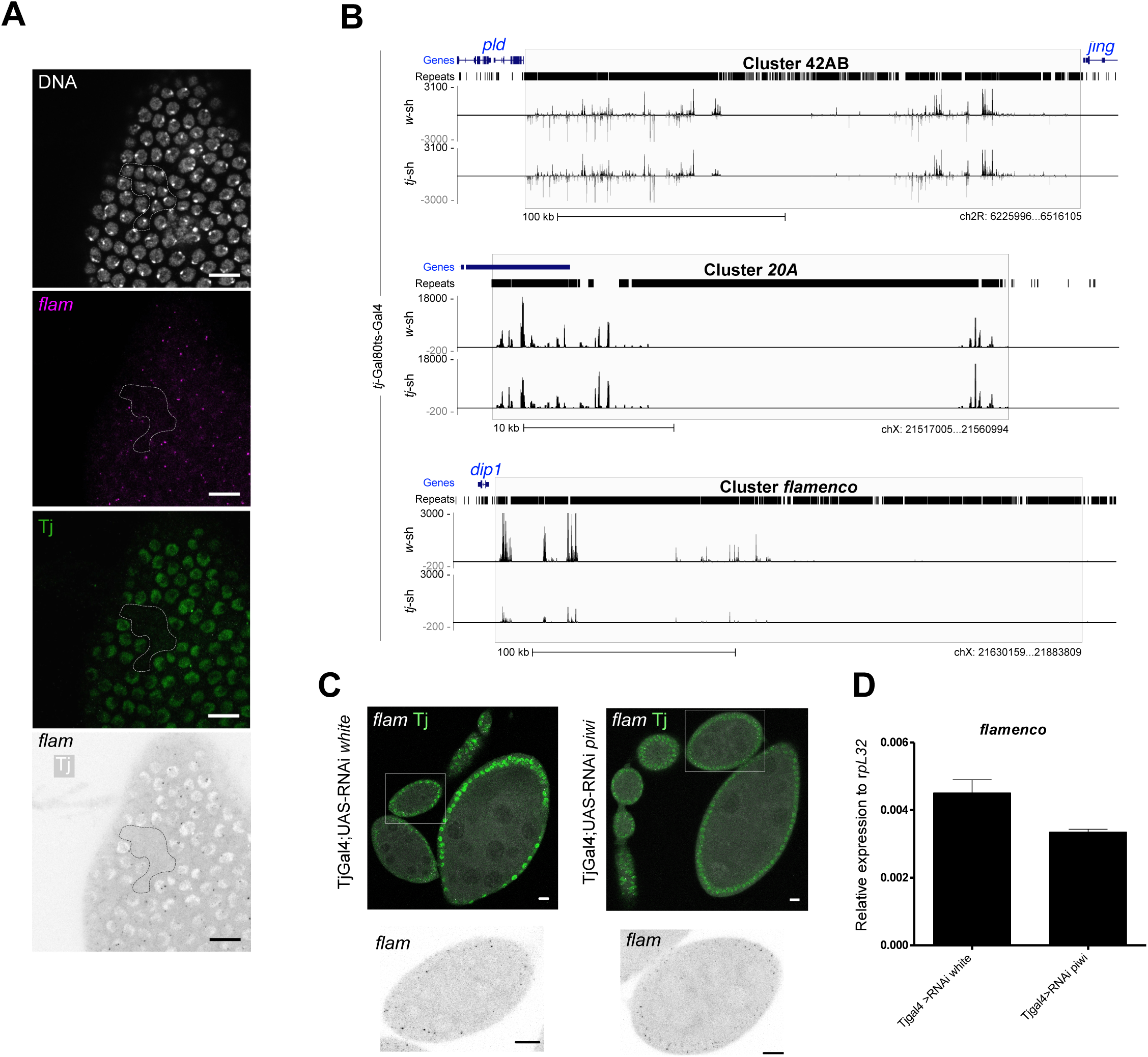
Tj controls *flam* expression. **(A)** Confocal images of *tj^eo^*^2^ clonal egg chamber showing *flam* RNA (magenta) by smFISH RNA and Tj (green) by immunostaining. The clone are circled in white. Nuclei are labelled with DAPI (white). Scale bars: 10 µm. **(B)** Uniquely mapped piRNAs are plotted over the *42AB*, *20A* and *flam* piRNA clusters. The piRNA profiles of *tj*-Gal80ts-Gal4; *w-sh* (control) and *tj*-Gal80ts-Gal4; *tj-sh* adult ovaries are shown. Normalized per million of unique genome-mapping piRNAs (0 mismatches). **(C)** Confocal images of egg chambers with indicated genotype showing *flam* RNA (magenta) and Tj (green) by Immuno-FISH. A zoom on one egg chamber (white square) is shown to the right. Scale bars: 10 µm. **(D)** Relative transgene expression in the indicate ovaries as measured by RT-qPCR. *tomato* expression levels were normalized to the expression of *rpL32.* Mean expression is shwon (n=6) with error bars representing standard deviation.

To investigate whether additional somatic transcription factors could be involved in the follicle cell-specific expression of *flam in vivo*, we carried out a Translating Ribosome Affinity Purification (TRAP) experiment. This method utilizes the UAS/Gal4 system and allows the identification of cell-specific messenger RNA (mRNA) translation^54^. In adult *Drosophila* ovaries, we applied TRAP to the GFP-labeled follicle cell population using the *tj*-Gal4 driver (UAS-GFP::Rpl10AGFP; *tj*-Gal4) and to the germ cell population with *nos*- Gal4 driver (UAS-GFP::Rpl10AGFP; *nos*-Gal4). Transcriptional profiling of purified mRNA followed by bioinformatic analyses identified around 900 genes with significantly higher expression and translation in somatic cells compared to germline cells (FC>2, p-value <0.05). In support of this, genes enriched in Tj positive cells were associated with specific gene ontology biological processes related to the development of follicle somatic cells (**Figure S4**). Among the differentially enriched genes, 30 were DNA binding factors and this list included Tj (**Table S2**). Using *tj*-Gal4 mediated somatic knockdowns (SKD), we next performed an RNAi screen targeting each candidate, and retained only those that reproduced the *flam* mutant phenotype that is characterized by atrophied ovaries^31,55^. We found that depletion of *tj*, *Stat92E*, *schnurri*, *tango*, *chd64* and *mamo* caused atrophic ovary development similar to *flam*. We next checked whether these factors affect *flam* expression by smFISH (**Figure S5**). Because SKD of *Stat92E*, *tango*, *chd64*, *mamo* and *schnurri* resulted in strongly atrophied ovaries, we instead transiently depleted these factors in adult follicle cells using a combination of *tj*-Gal4 the thermo-sensitive Tub-Gal80ts inhibitor^56^. A temperature shift from 18°C to 28°C enabled RNAi induction in late oogenesis stages, after ovaries had developed. In contrast to the atrophied ovary phenotype we observed with SKD, no impact on *flam* expression was detected following the depletion of *schnurri*, *tango*, *chd64*, and *mamo* using the thermo-sensitive RNAi approach (**Figure S5**). These results suggest that, while we cannot entirely rule out the contribution of multiple transcription factors, Tj appears to be the master TF responsible for soma-specific expression of *flam*.

### Selective derepression of *gypsy*-family TEs following Tj depletion

Since *flam* and the somatic piRNA pathway factors control *gypsy*-family TEs in OSCs, we next asked whether Tj depletion resulted in transposon derepression. RT-qPCR revealed ∼4-fold and ∼20-fold upregulation of *gypsy* expression following both 48 h and 96 h of *tj* siRNA treatment in OSCs (**Figures S6A and S6B**). RNA-seq from OSCs further confirmed robust upregulation of *gypsy* and other *gypsy*-family TEs after 48 h and 96 h of *tj* knockdown (**Figures 6A and S6C**). RT-qPCR from ovaries with *tj*-SKD showed upregulation of *gypsy* and *ZAM* (**Figure S6D**). Upregulation of *gypsy* itself was validated using a established reporter (*tj*-Gal4; *gypsy*-LacZ) (**Figure 6B**). Surprisingly, although several *gypsy*-family TEs were derepressed in *flam* mutant ovaries (**Figure 6C**), only a subset were upregulated in ovaries or OSCs depleted of Tj. The TEs *mdg1* and *412*, which are upregulated in *flam* mutants ovaries (**Figure 6C**), remained repressed upon *tj* knockdown in OSCs or ovaries (**Figures 6A and S6C**).

**Figure 6:**
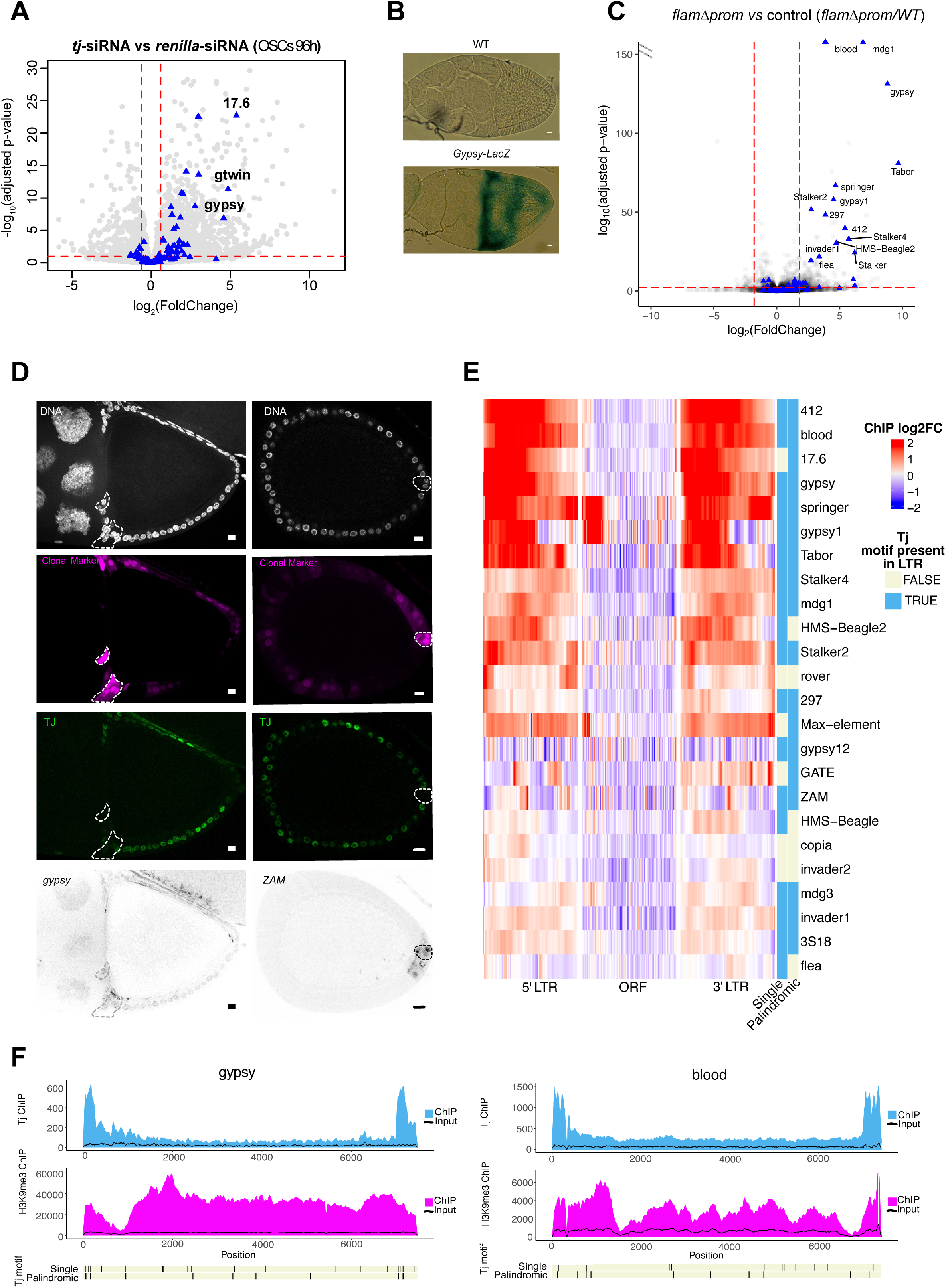
A subset of transposons appears directly regulated by Tj. (A) Volcano plot showing upregulation of *gypsy-*family TEs by 96 h of *tj* siRNA knockdowns in OSCs using differential mRNA-seq analysis (DEseq2) between *tj* and *renilla* siRNA knockdowns (mRNA-seq, n=3 replicates from distinct samples). Grey circles are showing the genes; blue triangles are showing the transposons. (B) Shown are beta-gal stainings of egg chambers as readout for gypsy silencing in *tj*-SKD and *w* SKD flies ovary (C) Volcano plot showing upregulation of *gypsy-*family TEs in *flamΔprom* flies ovary using differential mRNA-seq analysis between *flamΔprom*/*flamΔprom* and *flamΔprom/FM7* (DEseq2; n=3 replicates from distinct samples). Grey circles are showing the genes; blue triangles are showing the transposons. (D) Confocal images of *tj*-SKD clonal egg chambers showing *gypsy, ZAM RNA* (grey), Tj (green) and clonal marker (magenta) by Immuno-FISH. *tj*-SKD clones are circled in white. Nuclei are labelled with DAPI (white). Scale bars: 10 µm. (E) Heatmap showing log2 fold-change of Tj ChIP-seq signal over input for the indicated TEs that were selected based on expression in *flamΔprom* versus control fly ovaries or *tj* vs control knowckdown in OSCs (padj <0.01 and baseMean normalized by TE length in the top 70% of values for each dataset). (F) Position plots showing the ChIP-seq and input signals and positions of single and palindromic Tj motifs along the TE consensus sequence for the *gypsy* (top) and *blood* (bottom).

As *tj*-SKD ovaries were strongly atrophied, we validated our results in *tj*-SKD-induced mosaic clones. Consistent with our RT-qPCR, smFISH detected de-repression of *gypsy* and *ZAM*, but not *mdg1* or *412*, in *tj*-SKD-induced mosaic clones, compared to neighboring control cells (**Figures 6D and S6E**). We hypothesized that the differences in TE expression might be caused by some TEs also being directly regulated by Tj. Indeed, most of TEs deregulated in *flamΔprom* flies, such as *412* and *mdg1* showed strong Tj ChIP-seq peaks, particularly in the LTR region (**Figures 6E-F)**, which was distinct from H3K9me3 signal for the same TE. Co-opted regulation by Tj could potentially explain why certain TEs are not de-repressed following *tj* knockdown. In contrast, TEs such as *ZAM*, and *297,* which show little to no Tj binding in their promoter regions, are upregulated under both *flamΔprom* and *tj* knockdown conditions, consistent with their transcription being independent of the direct Tj binding. In addition, we noted that some TEs, such as *gypsy* display Tj binding yet remain de-repressed upon *tj*-SKD suggesting that other somatic transcription factors could contribute to their expression or that residual Tj protein could be sufficient for their activation. Of note, this is supported by the accompanying study by Rivera and colleagues which also showed that Tj binds and recognizes motifs within the LTRs of several TEs^57^. Overall, these results suggest co-evolution between Tj-dependent regulatory regions of TEs and piRNA pathway genes and clusters.

## Discussion

Our study identifies Tj as a pivotal transcriptional regulator of the somatic piRNA pathway in *Drosophila* ovaries and adds a key layer to our understanding of TE repression mechanisms in *Drosophila*, particularly in non-germline tissues. Previous studies have demonstrated Tj’s involvement in gonadal development and its regulation of *piwi* expression^33,34^. Our findings expand on these roles by demonstrating Tj’s broader control over somatic piRNA pathway components in ovarian follicle cells. Our results reveal Tj as a regulator of somatic piRNA pathway components, including *fs(1)Yb*, *nxf2*, *panx*, and *armi*. ChIP-seq and motif analyses demonstrate that Tj binding at conserved promoter motifs is critical for the expression of these genes. Moreover, Tj regulates the soma-expressed piRNA cluster *flam*, which is essential for TE silencing. This dual regulation of piRNA factors and clusters mirrors the function of mammalian transcription factors such as A-MYB, TCFL5 and BTBD18 which coordinate piRNA pathway gene expression and pachytene piRNA cluster activation during spermatogenesis^58,59^.

Beyond its role in piRNA regulation, Tj appears to be a master regulator of somatic cell identity with approximately 8,000 Tj binding sites detected at the promoters of somatically enriched genes. RNA-seq analyses of *tj* knockdowns in OSCs revealed a significant shift from somatic to germline gene expression, with upregulation of germline-specific genes such as *osk*, *ovo*, and *nos* (**Table S1**). This shift is accompanied by the downregulation of *l(3)mbt*, suggesting that Tj maintains the somatic expression program by repressing germline gene transcription. Interestingly, Tj and Ovo, a germline-specific transcription factor^38^, exhibit opposing binding patterns at the promoters of somatic and germline genes (**Figure S7A**), underscoring their complementary roles in maintaining lineage-specific gene expression programmes within gonads (**Figure S7B**). However, while Ovo’s role in regulating germline piRNA factors is conserved across species, it is not clear whether Tj’s regulation of the somatic piRNA pathway is a *Drosophila*-specific adaptation or more broadly conserved. We speculate that this evolutionary innovation likely arose in response to the challenge posed by *gypsy*-family TEs, which can infect the germline via somatic expression^60,61^. These TEs have evolved distinct expression patterns in somatic follicle cells^60,61^. When the piRNA pathway is disrupted, some TEs are broadly expressed, while others remain restricted to specific niches, *ZAM* in posterior follicle cells and *Idefix* in the germarium for example^60–62^, indicating regulation by distinct transcription factors. Indeed, some TEs were reported to rely on specialized transcription factors, such as the Pointed for *ZAM*^63^, while our results indicate that other TEs are likely co-regulated by Tj, allowing for broader or more robust somatic expression. Notably, only a subset of TEs derepressed in *flam* mutants are also affected by Tj loss, suggesting that some TEs are independent of Tj for their expression. Co-evolution between TEs and the piRNA pathway could suggest an ongoing evolutionary “arms race” between host regulatory mechanisms and TEs. The coordinated expression of *flam* and somatic piRNA pathway components likely reflects the selective pressures exerted by TEs that shaped their *cis*-regulatory sequences to respond to one key transcription factor in ovarian somatic cells allowing for efficient and coordinated silencing of both niche-specific and broadly expressed TEs in somatic cells.

Although primarily considered a transcriptional activator, Tj may also function as a repressor - either at the transcriptional level by silencing adhesion molecule genes or post-transcriptionally through its 3’ UTR, which generates piRNAs that target and silence specific genes such as *FasIII*^33,34^. The precise mechanisms by which Tj regulates its targets remain to be fully elucidated, particularly the potential involvement of additional transcriptional regulators and co-factors. Other factors, such as Ci, Mirror or Stat92E, which are all DNA-binding proteins expressed in somatic follicle cells that were identified in our TRAP-based screening, could contribute to the regulation of piRNA genes and *flam*, as is often described in gene regulatory networks where multiple transcription factors work together to ensure robustness and precise control across development^64,65^.

Tj’s similarity to mammalian Maf proteins, c-Maf and MafB in particular, including its potential for dimerization, adds another layer of specificity to its regulatory function. Like other members of this TF family, Tj contains a leucine-zipper domain, a basic DNA-binding domain, and a Maf-specific extended homology domain in its C-terminal region.

The leucine-zipper domain, known for facilitating the dimerization of Maf proteins, either with other Maf or bZIP TFs (e.g., homo/heterodimeric AP-1 complexes), suggests that Tj could also function as a dimer^33^. The presence of pseudo-palindromic motifs within Tj binding sites, similar to the MARE motifs recognized by the dimeric Maf family proteins, further reinforces this idea^40,41^. Interestingly, the pseudo-palindromic Tj motifs at promoters of the somatic piRNA factors were often mutated (**Figure S1B**), which has been previously shown to confer DNA-binding specificity for homodimeric bZip transcription factor complexes over their heterodimeric versions in EMSA experiments^41^.

In conclusion, our findings establish Tj as a master regulator of the somatic piRNA pathway and a key player in maintaining genome integrity through TE repression. By regulating both piRNA pathway components and the *flam* piRNA cluster, Tj exemplifies the efficiency of a single transcription factor in orchestrating complex gene networks. Tj’s interaction partners which likely contribute to its finely tuned regulatory expression network, await further exploration.

## Limitations of the Study

While our data suggest direct effects of Tj on the somatic piRNA pathway, we cannot entirely exclude the possibility of indirect effects. Tj regulates numerous somatic genes, including transcription factors, and its broad regulatory network may contribute indirectly to piRNA pathway activation. For example, the downregulation of *l(3)mbt*, a repressor of germline programs in somatic cells, observed upon *tj* knockdown could indirectly influence somatic piRNA pathway gene expression. Further investigation will be needed to disentangle these direct and indirect regulatory effects. Tj is known to regulate many genes critical for ovarian development, and therefore the ovarian atrophy observed upon Tj loss could be a consequence of both TE reactivation due to piRNA pathway disruption and/or misregulation of developmental genes critical for ovarian tissue homeostasis. In addition, our results with the *de novo* motif analysis with STREME did not yield the pseudo-palindromic motifs likely due to limitation in the detection of gapped motifs by the software, as well as due to frequent mutations and less frequent appearance of the pseudo-palindromes at specific gapped lengths (1-4bp) within the peaks; however, we were able to map them using the motif enrichment (i.e., CentriMo in MEME-ChIP) and scanning tools (i.e., FIMO).

## Author Contributions

AzA performed experiments and bioinformatic analyses, designed figures, and contributed to manuscript writing. AM conducted data analysis, Tj clonal experiments, functional assays with transcription factor knockdown, prepared figures, and contributed to manuscript writing. NM performed all the cloning, generated *Drosophila* transgenic lines, and conducted Tomato immunostaining and corresponding small RNA sequencing. JRS performed the experiments, analyzed, plotted RT-qPCR data and contributed to manuscript writing. BB designed CRISPR constructs, generated the corresponding *Drosophila* mutant lines, conducted TRAP experiments and bioinformatics analyses. NG was responsible for smFISH probe labeling. VM conducted TRAP experiments and bioinformatics analyses. AMP performed computational analyses. NCL designed the Tj peptide, funded antibody production, and supervised AJR in conducting affinity purification and validating the specificity of the anti-Tj antibody. AbA contributed to the Tj clonal experiments and performed ovary RT-qPCR experiment, analyzed and plotted the data. SMM contributed to the Tj clonal experiments, analysis, and manuscript writing. GJH supervised the project and contributed to funding acquisition. BCN conceptualized the project, designed the experiments, supervised the work, contributed to funding acquisition and wrote the manuscript. EB conceptualized the project, designed the experiments, supervised the work, contributed to funding acquisition and wrote the manuscript.

## Acknowledgments

We thank Brasset and Hannon group members for fruitful discussions, and I. Vaillant for comments on the manuscript and S. Jensen for participating in the bioinformatics training of students. We thank the Bloomington *Drosophila* Stock Center for providing fly lines. We thank Dorothea Godt for providing Tj mutant *Drosophila* stocks. We thank Y. Renaud and P. Pouchin for helpful advices and development of the iGReD bioinfomatics platform. We also thank the CLIC facility (Clermont Imagerie Confocale). We thank the Scientific Computing core at the CRUK Cambridge Institute for HPC resources, Research Instrumentation and Cell Services for training, liquid nitrogen storage and mycoplasma testing, and the Genomics core for sequencing services. This work was supported by grants from the Agence Nationale pour la Recherche (CHApiTRE ANR-20-CE12-0005, EB, BiopiC ANR-21-CE12-0022, EB). AM was supported by the Fondation pour la Recherche Médicale (ECO202006011583 and FDT202304016393, AM). This research was financed by the French government IDEX-ISITE initiative 16-IDEX-0001 (CAP 20-25). NCL is funded by NIH/NIGMS grant R01GM135215. GJH is a Royal Society Wolfson Research Professor (RSRP\R\200001). This research was funded in whole, or in part, by Cancer Research UK (G101107) and the Wellcome Trust (110161/Z/15/Z and 226627/Z/22/Z).

## Supplementary Figure legends

**Figure S1:**
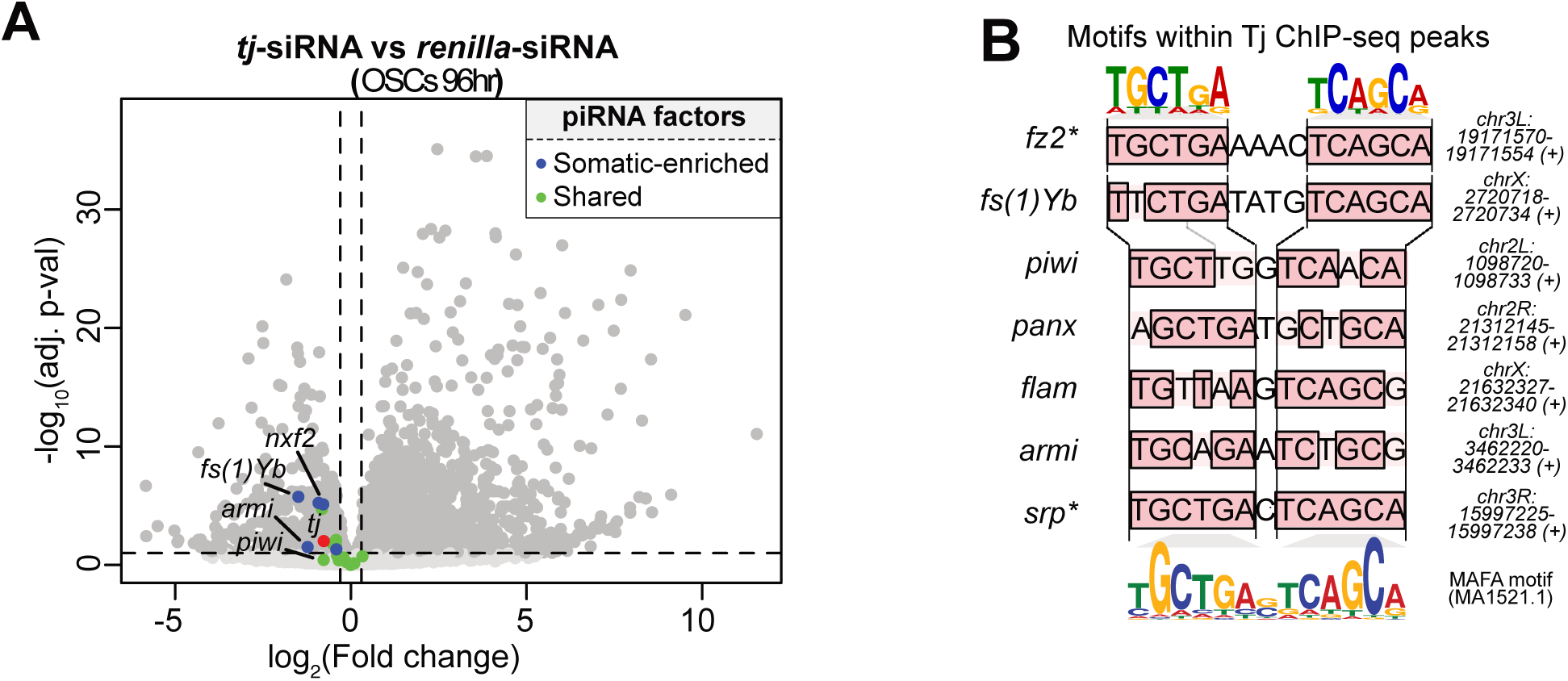
Knockdown of *tj* in OSCs downregulates somatic piRNA pathway genes. **(A)** Volcano plot showing differential RNA-seq analysis (DEseq2) between *tj* and *renilla* siRNA knockdowns (96 h; n=3 replicates from distinct samples) in OSCs. Blue dots are showing the soma-enriched piRNA pathway genes (*fs(1)Yb*, *nxf2*, *panx*, *soYb* and *armi*); green dots showing general piRNA factors (e.g., *piwi*); red dots showing *tj*. **(B)** Pseudo-palindromic Tj motifs are found within Tj ChIP-seq peaks at promoters of somatic piRNA pathway components and somatic non-piRNA pathway target genes (indicated by asterisk). Motif logos shown above were generated from the shown Tj motif half-sites. Motif logo for MAFA shown below was taken from JASPAR database (MA1521.1).

**Figure S2:**
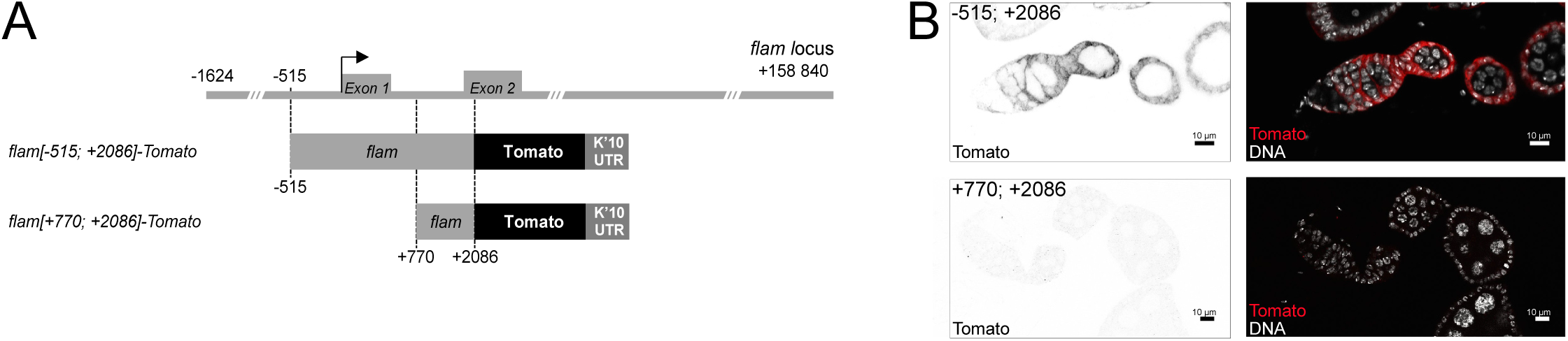
The +770; +2086 region of *flam* is insufficient for its transcription. **(A)** Schematic representation of the transgenic constructs. **(B)** Confocal images of egg chambers with the indicated genotypes showing Tomato expression (red) by immunostaining. Nuclei are stained with DAPI (white). Scale bars: 10 µm.

**Figure S3:**
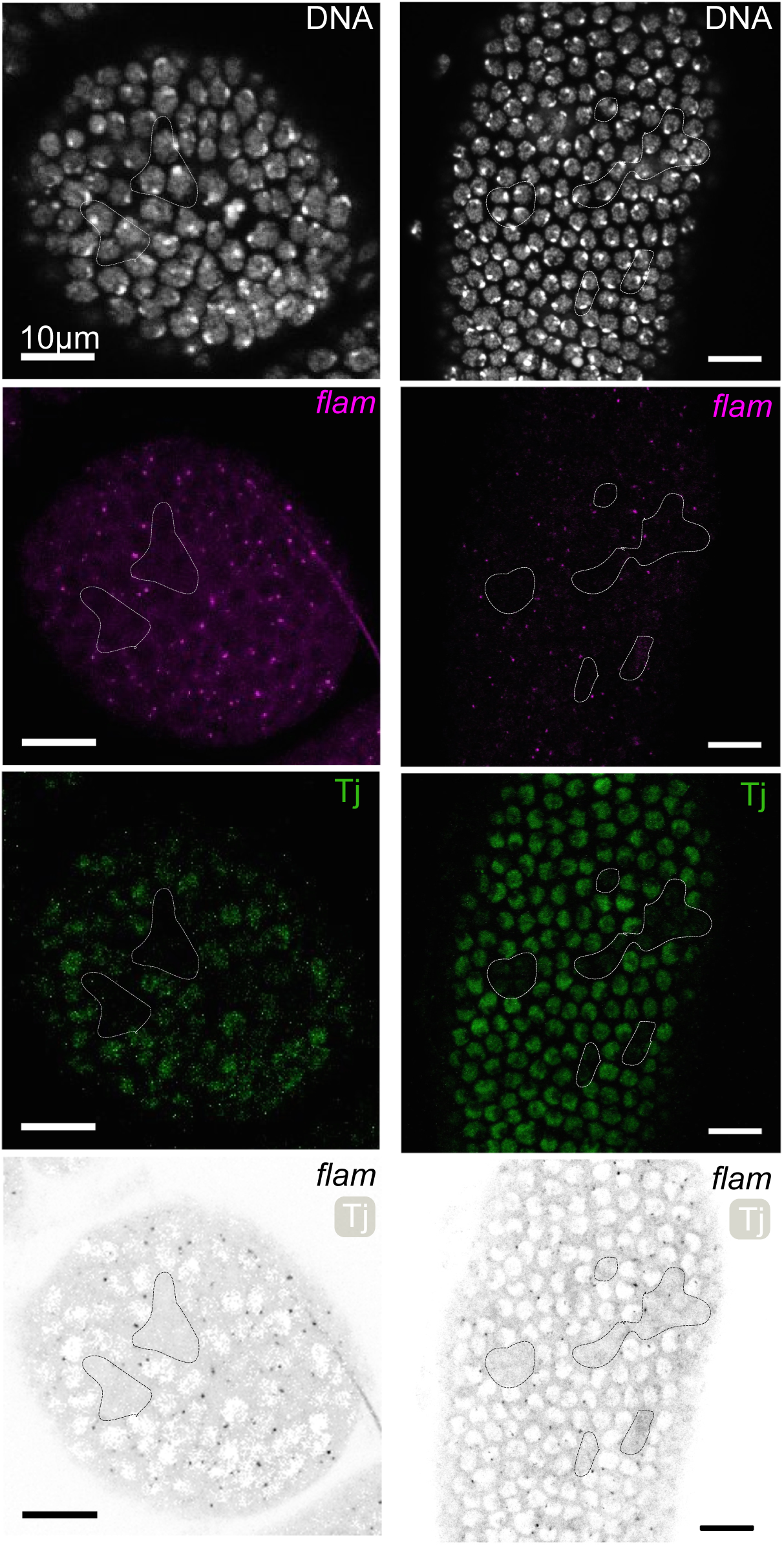
Tj depletion disrupts *flam* expression. Confocal images of *tj^eo2^* clonal egg chambers showing *flam* RNA (magenta) and Tj protein (green) levels measured by Immuno-FISH. *tj^eo2^*clones are circled in white. Nuclei are labelled with DAPI (white). Scale bars: 10 µm.

**Figure S4:**
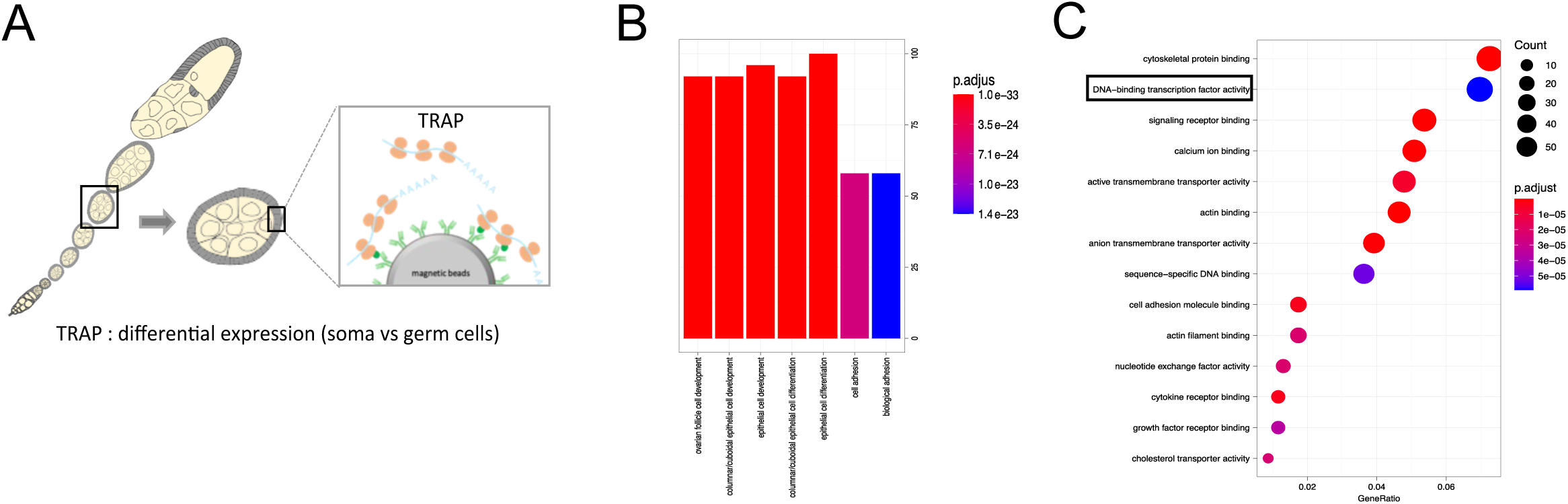
TRAP experiments identify additional candidate TFs regulating *flam* expression. **(A)** Schematic representation of the Translating ribosome affinity purification (TRAP) method performed on somatic or germ cells of ovaries expressing GFP-tagged ribosomal protein RpL10a, driven by *tj-*Gal4 or *nos*-Gal4 respectively. Comparison between UAS-GFP::Rpl10AGFP; *tj*-Gal4 to UAS-GFP::Rpl10AGFP; *nos*-Gal4 conditions. **(B)-(C)**. Gene ontology analyses of genes enriched in follicles cells. List of the significantly enriched GO Biological Process **(B)** and Molecular Function **(C)** terms.

**Figure S5:**
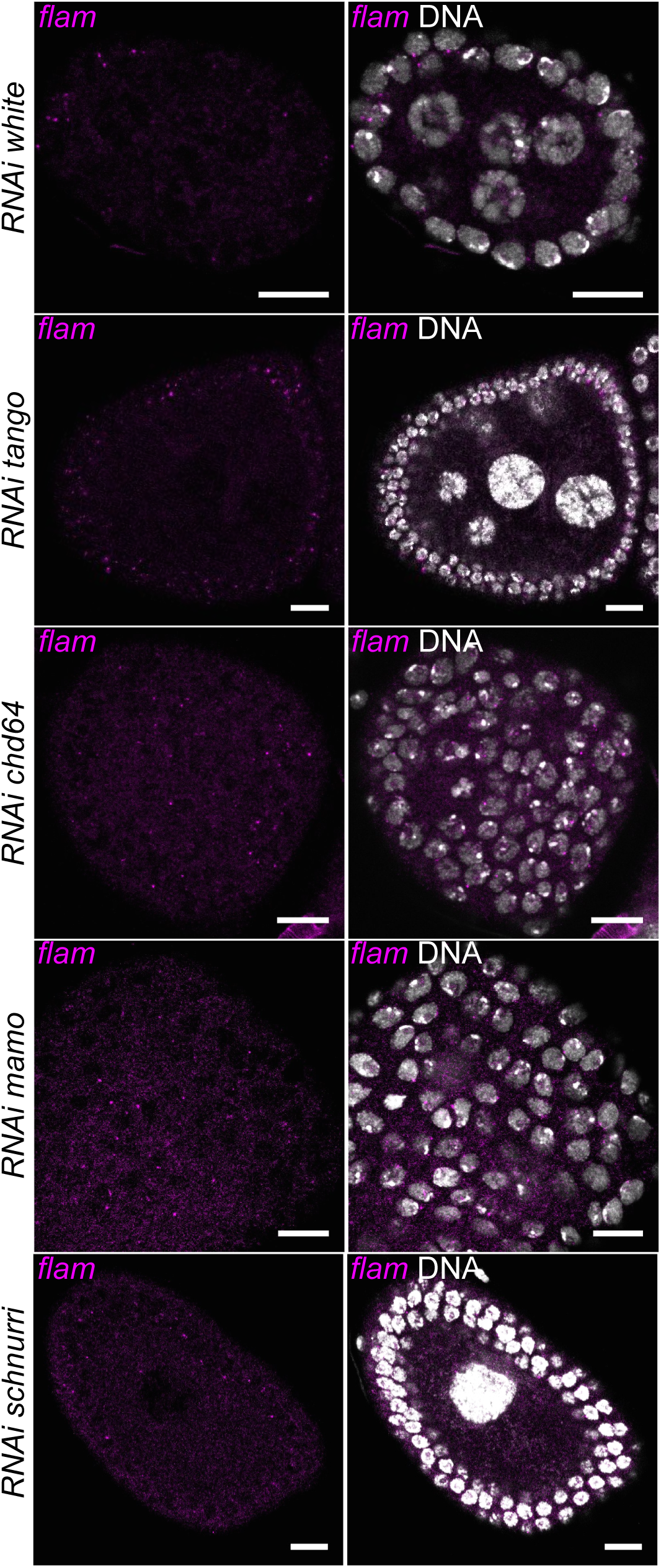
Effects of candidates from TRAP experiment on *flam* expression. Confocal images of egg chambers with indicated genotype stained for *flam* RNA using smRNA-FISH (magenta). The driver used is *tj*-Gal80ts-Gal4. Nuclei are labelled with DAPI (white). Scale bars: 10 µm.

**Figure S6.**
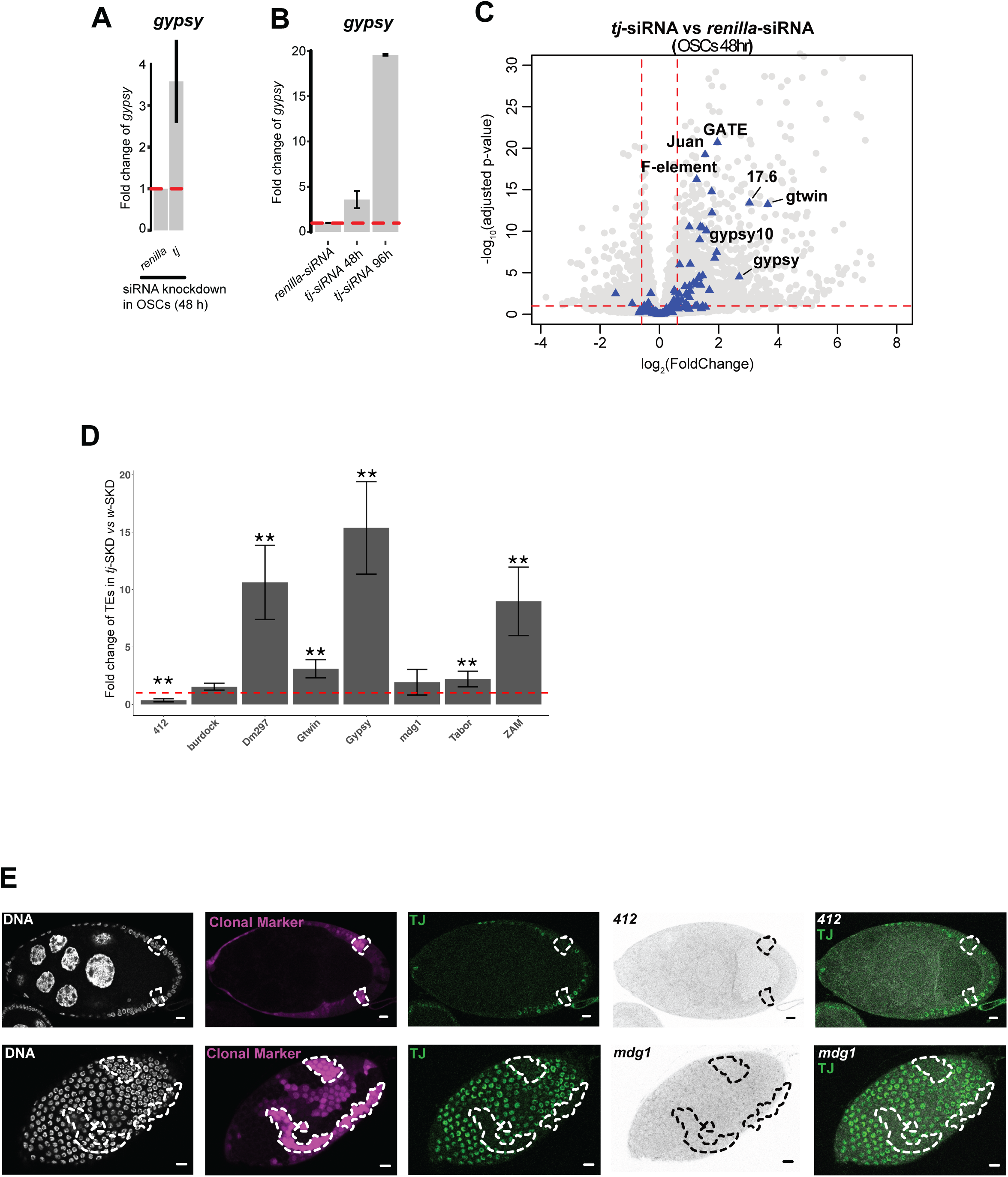
: A subset of transposons appears directly regulated by Tj. **(A)** Fold-changes in *gypsy* TE expression upon *tj* knockdown in OSCs after 48 h are shown by RT-qPCR (*rpL32* housekeeping control, n=3, error bars indicate standard deviation). **(B)** As in **(A)** but comparing fold-changes between 48 h and 96 h of *tj* knockdown **(C)** Volcano plot showing upregulation of transposons by 48 h of *tj* siRNA in OSCs using differential mRNA-seq analysis (DEseq2) compared to *renilla* siRNA knockdowns (mRNA-seq, n=3 replicates from distinct samples). Grey circles are showing the genes; blue triangles are showing the transposons. **(D)** Fold-changes in TE expression upon *tj*-SKD in ovaries are shown by RT-qPCR (*rpL32* housekeeping control, n=3, error bars indicate standard deviation) **(E)** Confocal images of *tj*-SKD clonal egg chambers showing *mdg1, 412 RNA* (grey), Tj (green) and clonal marker (magenta) by Immuno-FISH. *tj*-SKD clones are circled in white. Nuclei are labelled with DAPI (white). Scale bars: 10 µm.

**Figure S7:**
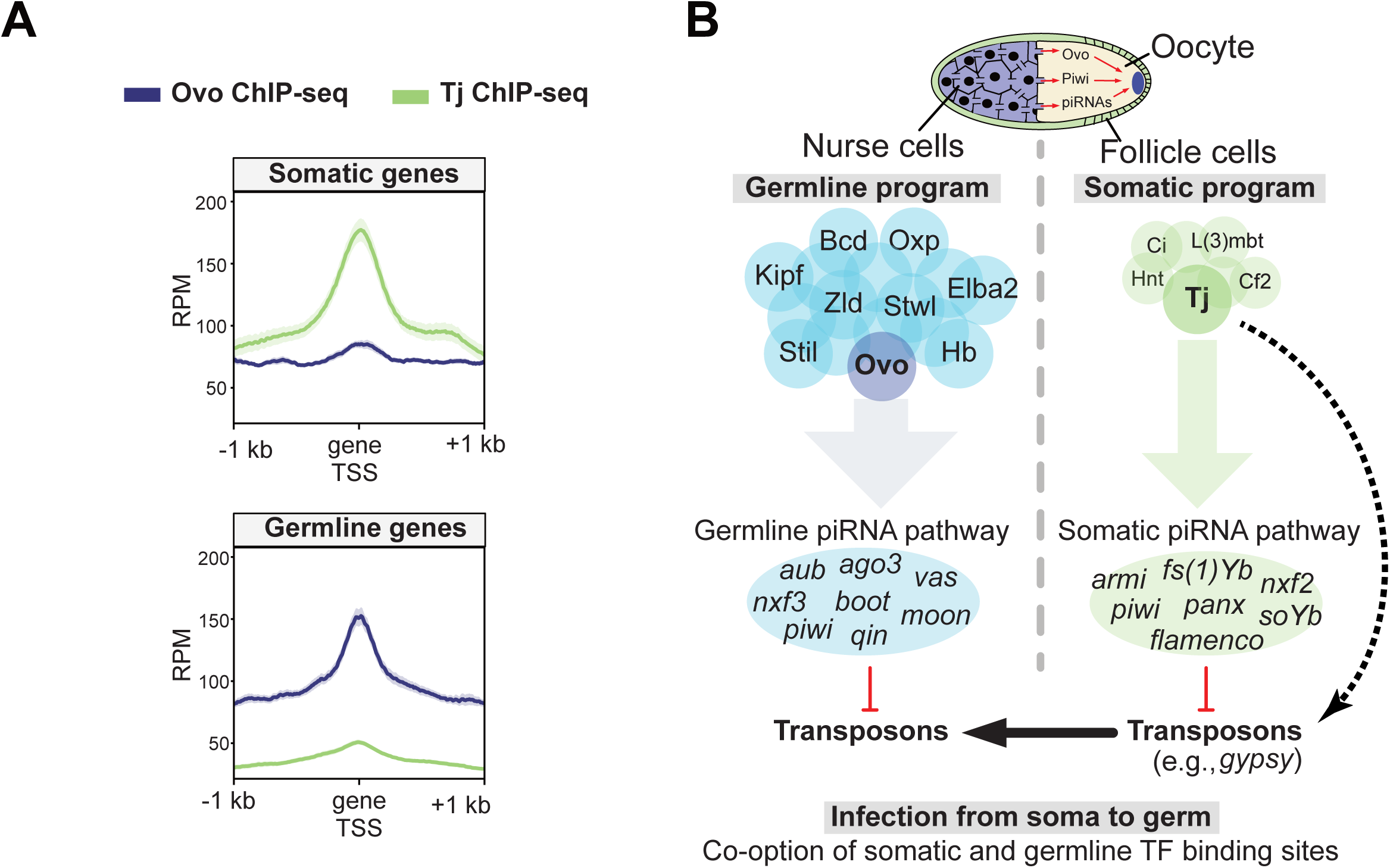
Control of germline and somatic piRNA pathway components relies on Ovo and Tj. **(A)** Ovo and Tj ChIP-seq showing antagonistic binding at the promoters of germline and somatic genes. **(B)** Model depicting the germline and somatic transcriptional regulators of the piRNA pathway components in *Drosophila* ovaries and their putative co-option by transposons for mobilization and infection from soma into the germline.

## STAR★Methods

### Key resources table

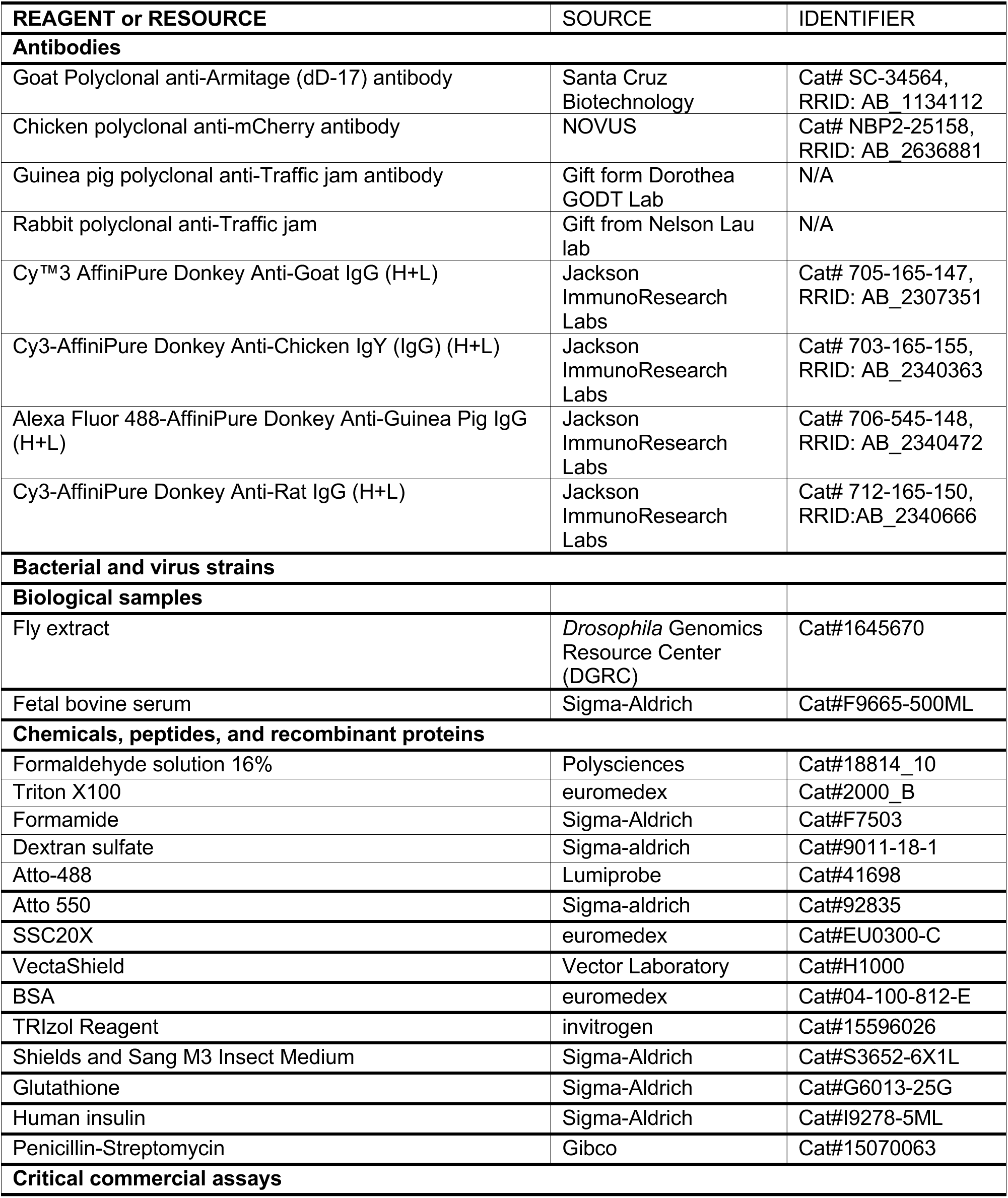
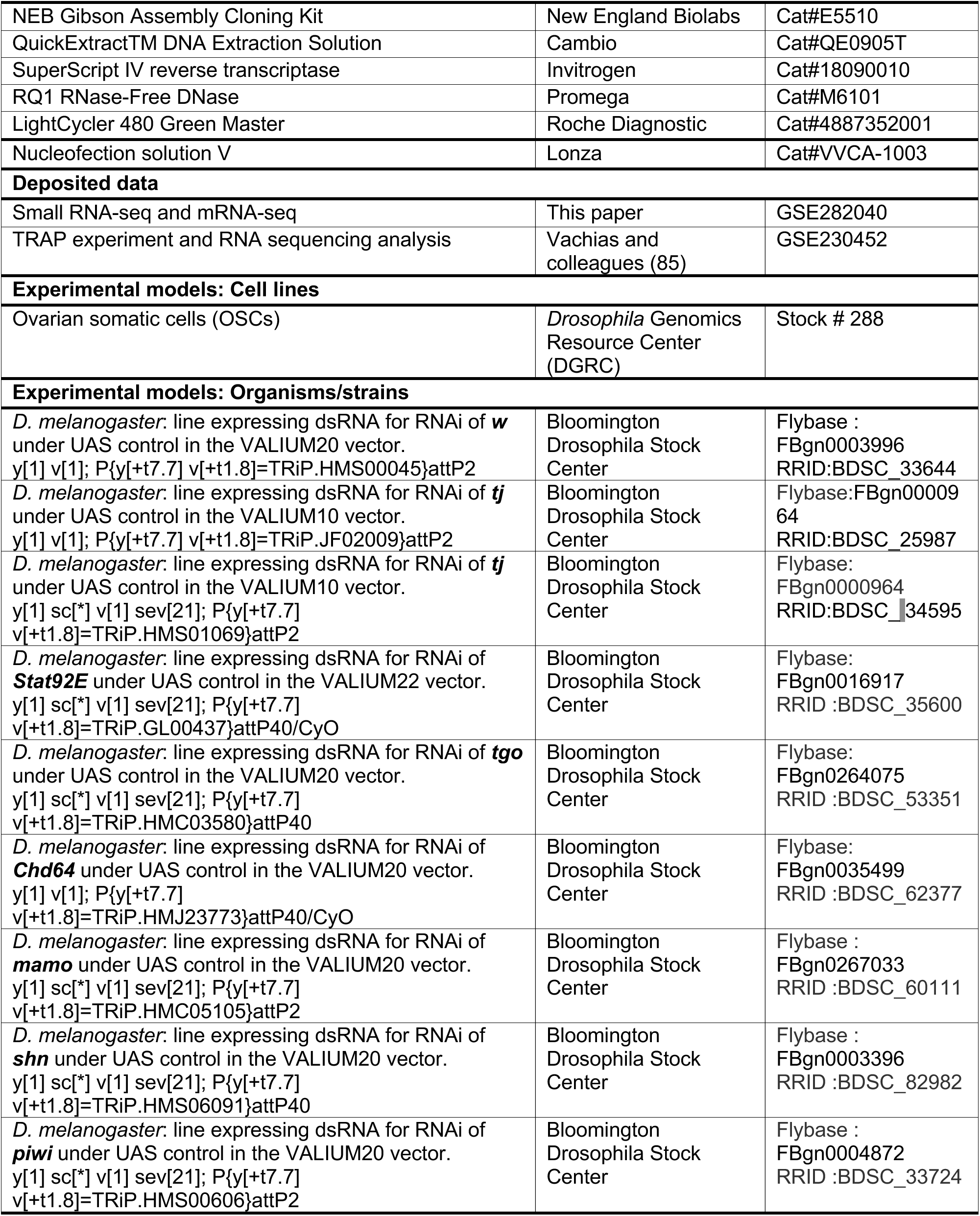
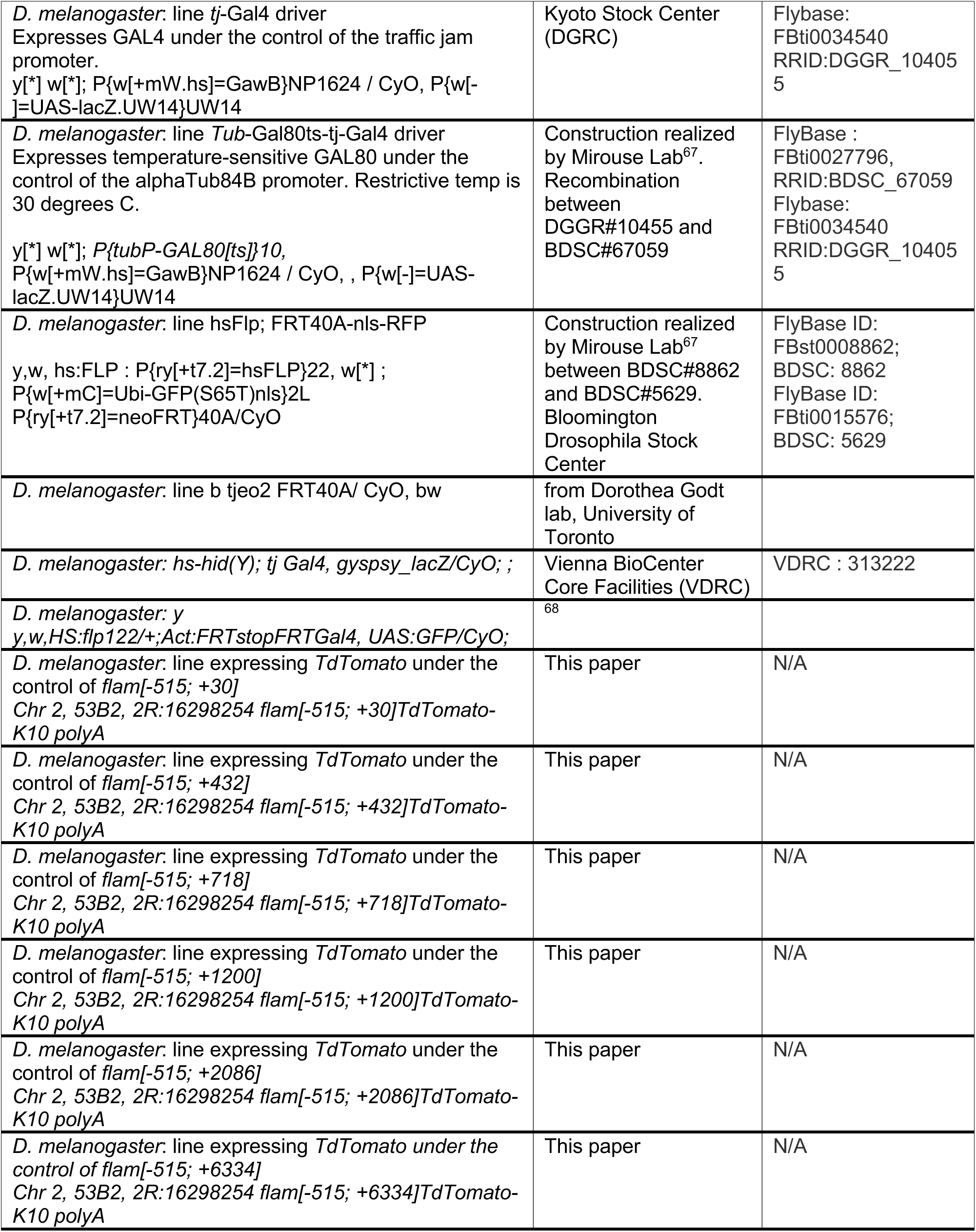
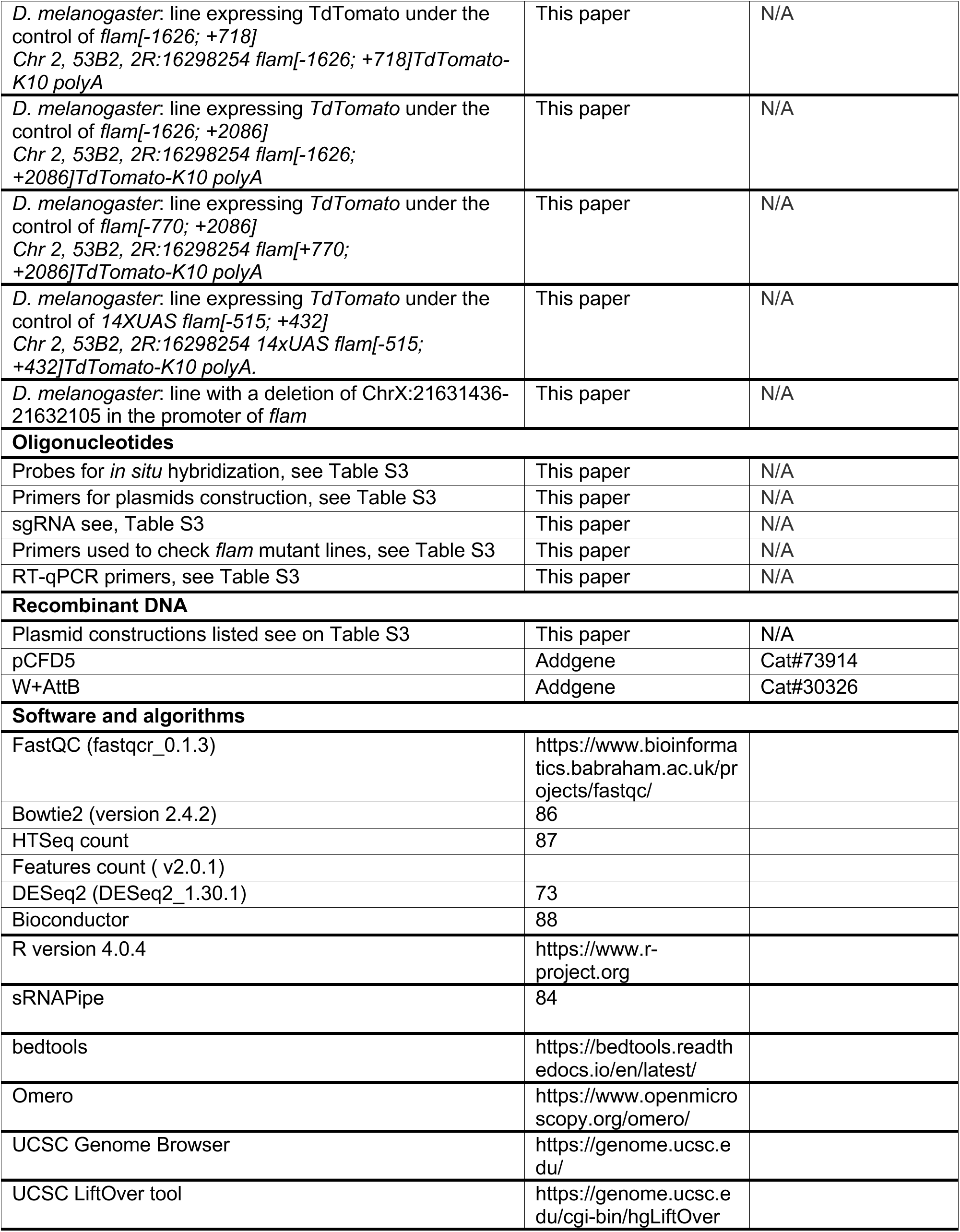
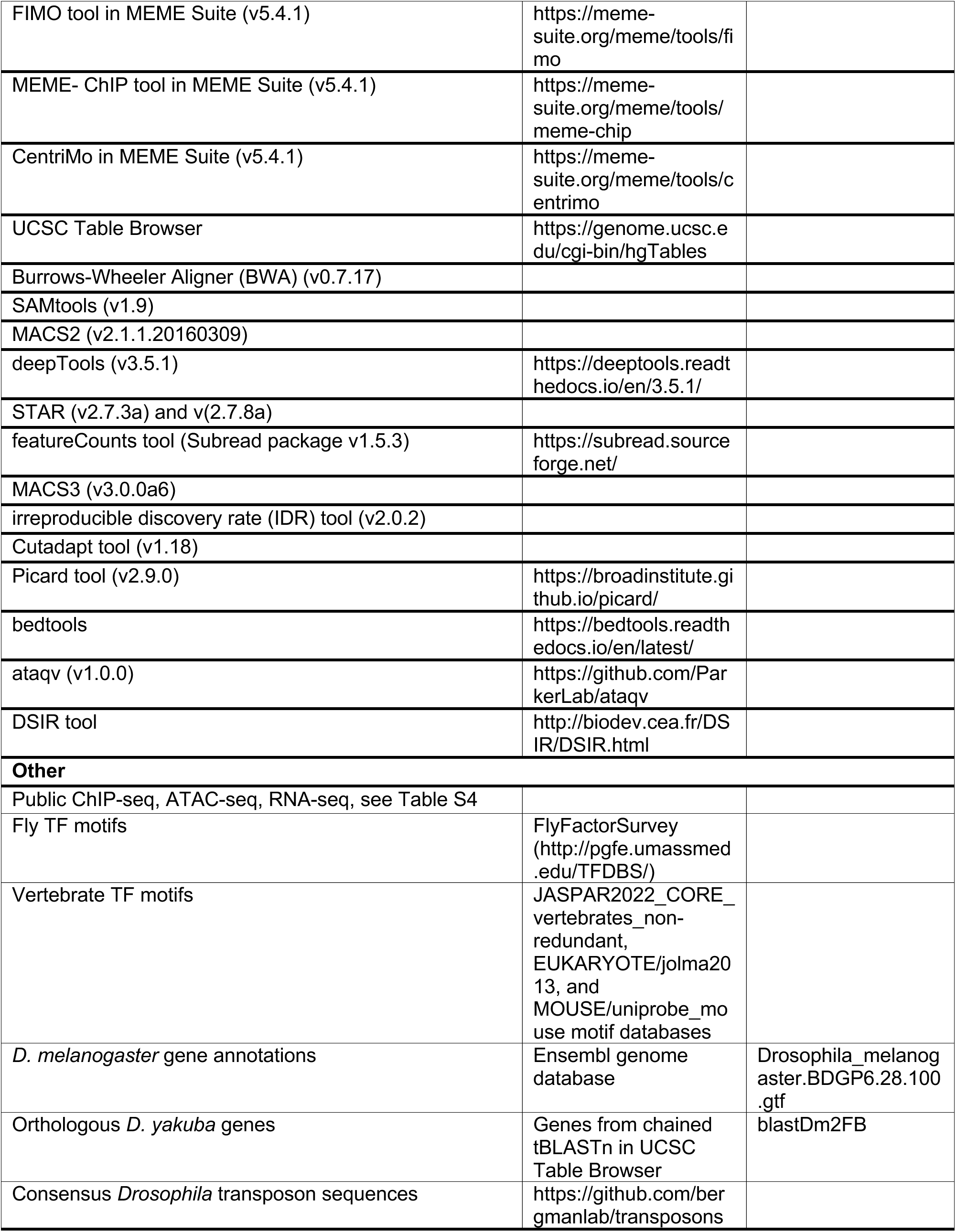

### Experimental model and study participant details

#### D. melanogaster flies

*Genotypes: listed in Key resources table. Age: Adult stages, 3 to 6 days old*

*Sex: female (to study ovaries)*

*Housing: 20°C or 25°C, on standard medium.*

*For heat shock experiments flies are placed 1h at 37°C on standard Medium.*

#### Wild-type OSCs

*Species: D. melanogaster*

*Sex: female*

*Tissue: ovary*

*Growth media: cultured at 26 °C in Shields and Sang M3 Insect Medium (Sigma-Aldrich, S3652-6X1L) supplemented with 10% fetal bovine serum (Sigma-Aldrich, F9665-500ML), 10% fly extract (DGRC, Stock # 1645670), 0.6 mg/mL glutathione (Sigma-Aldrich, G6013-25G), and 10 mU/mL human insulin (Sigma-Aldrich, I9278-5ML) + 1% Penicillin-Streptomycin (Gibco, 15070063).*

### Method details

#### Cell culture and growth media

Wild-type OSCs^34^ were purchased from the *Drosophila* Genomics Resource Center (DGRC, cat # 288) and cultured at 26°C in Shields and Sang M3 Insect Medium (Sigma-Aldrich, cat # S3652-6X1L) supplemented with 10% fetal bovine serum (Sigma-Aldrich, cat # F9665-500ML), 10% fly extract (DGRC, cat # 1645670), 0.6 mg/mL glutathione (Sigma-Aldrich, cat # G6013-25G), 10 mU/mL human insulin (Sigma-Aldrich, cat # I9278-5ML), and 1% Penicillin-Streptomycin (Gibco, cat # 15070063).

#### siRNA knockdowns in cell culture

Sense and antisense 21 nt siRNA sequences for target genes were designed using the DSIR tool (http://biodev.cea.fr/DSIR/DSIR.html) (**Table S3**) and ordered from IDT. The RNA oligonucleotides were resuspended to 400 μM final concentration in RNase-free water. Equal volumes of the resuspended sense and antisense siRNA were mixed and then added to an equal amount of 2x annealing buffer (60 mM potassium acetate; 200 mM HEPES, pH 7.5). The mix was boiled for 5 min at 75°C, then ramped down to 25°C (-0.1°C/s) to anneal the siRNA sequences into the siRNA duplex (100 μM final concentration). 2 μL of the final 100 μM siRNA duplex was mixed with 100 μL of Nucleofection solution V (Lonza, VVCA-1003) and transfected into 4 million cells (48 h knockdown) or 5 μL of 100 μM siRNA duplex was mixed with 100 μL of Nucleofection solution V and transfected into 10 million cells (96 h knockdown) using a NucleofectorTM 2b Device and program T-029 (Lonza). Cells were plated into 6-well plates (5 million cells) or 10 cm plates (10 million cells). RNA was either harvested at 48 h or nucleofection was repeated after 48 h and RNA was harvested at 96 h. RNA was either harvested at 48 h or nucleofection was repeated after 48 h and RNA was harvested at 96 h.

#### RNA isolation and RT-qPCR

OSCs were homogenized by addition of Buffer RLT directly on the cell culture vessel surface. RNA was isolated using RNeasy Mini Kit (QIAGEN, 74106) with RNase-Free DNase Set (QIAGEN, 79254) for DNase digestion during RNA purification as per manufacturer’s instruction. Reverse transcription was performed with 100 ng to 1 μg RNA using SuperScript IV Reverse Transcriptase (Thermo Fisher Scientific, 18090010). RT-qPCR was performed on 1:10 diluted cDNA using Fast SYBR Green Master Mix (Thermo Fisher Scientific, 4385610) with Bio-Rad C1000 Thermal Cycler. Primers were designed against exon-exon junctions for genes with introns and *rpL32* was used as an internal standard (**Table S3**). Relative expression was analysed using the delta-delta Ct method^62^. When total RNAs were extracted from fresh ovaries, we used TRIzol (Invitrogen) following the manufacturer’s instructions. After DNase treatment (RQ1 RNase-free DNase Promega), cDNA was synthesized using random priming of 1 μg total RNA and SuperScript IV Reverse Transcriptase (Invitrogen). Quantitative PCR was performed with the LightCycler480 SYBR Green I Master mix system (Roche), and the data were analyzed using the LightCycler480 system (Roche). Each experiment was performed 6 times with technical duplicates. RNA levels were calculated with the relative quantification method with *rpL32* housekeeping gene as the reference (for primer sequences, see **Table S3**). Statistical differences were evaluated using Kruskal-Wallis non-parametric test, with corrections for multiple comparisons (Holm–Bonferroni adjustment method).

For mRNA-seq experiments, total RNA was extracted from 30 ovaries using TRIzol (Invitrogen) following the manufacturer’s instructions.

#### RNA-seq

For mRNA-seq from OSCs, selection was performed using Poly(A) mRNA Magnetic Isolation Module (NEB, E7490) with 1 ug of total RNA as input. Libraries were prepared using The NEBNext Ultra Directional RNA Library Prep Kit for Illumina (NEB, E7420L). Library quality was assessed using Agilent 2100 bioanalyzer (High Sensitivity DNA Kit). Libraries were quantified with KAPA Library Quantification Kit for Illumina (Kapa Biosystems, cat# KK4873) submitted to sequencing on a NovaSeq X System with 2 × 50 bp reads.

For mRNA-seq from ovaries 1 µg of total RNA was send to BGI for library construction using the DNBSEQ Eukaryotic Strand-specific mRNA library workflow. Libraries were sequenced on platform DNBSEQ to a read length of 150 bp paired-end. Raw data were filtered by BGI in order to remove adapter sequences or low-quality reads.

#### Fly stocks and crosses

All *D. melanogaster* stocks were grown on standard medium at 20°C. RNAi lines expressing dsRNAs^80^ or shRNAs^81^ against *w* (#33644), *tj* (#25987, #34595), *Stat92E* (#35600), *tango* (#53351), *chd64* (#62377), *mamo* (#60111)*, schnurri (#82982), piwi* (#33724) were obtained from the Bloomington *Drosophila* stock center (BDSC). The *tj*-Gal4 driver was originated from Kyoto Stock Center DGRC, #104055. The *gypsy*-lacZ strain (#313222) was obtained from Vienna *Drosophila* Resource Center (VDRC). The lines *tub-*Gal80ts-tj-Gal4 driver and hsFlp; FRT40A-nls-RFP were a kind gift of V. Mirouse at iGReD^67^ and the *tj^eo^*^2^ FRT40A/CyO, bw line from Dorothea Godt (University of Toronto). The line *y,w,HS:flp122/+;Act:FRTstopFRTGal4, UAS:GFP/CyO;* is a kind gift of C. De Joussineau. The *tj*-Gal4 driver strain was crossed with the transgenic 14xUAS *flam*[-515; +482] *tomato* flies and with all RNAi strains. *tj-*Gal80ts-Gal4 line was crossed with RNAi lines except RNAi lines against *piwi*; the Gal80ts and Gal4 are driving by *tj*; Gal80ts is inactivated at 30°C. *tj*-Gal4 driver was crossed with shRNA *tj* or shRNA *w* as a control at 25°C. F1 flies obtained were used for mRNA-seq analysis.

#### Clones and Constructs

All plasmids were constructed using NEB Gibson Assembly Cloning Kit (cat # E5510) in accordance with the manufacturer’s instructions and the recommendations of the NEBuilder assembly tool (https://nebuilder.neb.com/). Inserts (listed in the **Table S3**) were amplified by PCR using Phusion taq polymerase (Thermofisher) using primers summarized in **Table S3**, and cloned in the W+attB plasmid (addgene #30326). Ten vectors have been then elaborated in W+attB backbone: nine of them possessed different *flam* fragments sizes upstream of tdTomato reporter sequence directly followed by polyA signal sequence from *K10* gene:

*flam[-515; +30]TdTomato-K10 polyA*, *flam[-515; +432]TdTomato-K10 polyA*, *flam[-515;*

*+718]TdTomato-K10 polyA*, *flam[-515; +1200]TdTomato-K10 polyA*, *flam[-515;*

*+2086]TdTomato-K10 polyA*, *flam[-515; +6334]TdTomato-K10 polyA*, *flam[-1626;*

*+718]TdTomato-K10 polyA*, *flam[-1626; +2086]TdTomato-K10 polyA*, *flam[+770;*

*+2086]TdTomato-K10 polyA*. The last construct contains the fourteen UAS sequence upstream of the *flam[-515; +432]TdTomato-K10 polyA fragment and is named 14xUAS flam[-515; +432]TdTomato-K10 polyA*.

#### Transgenic animal production

Plasmid constructions were purified using Plasmid Plus Maxi Kit (QIAGEN). Purified DNA was used for PhiC31 integrase-mediated transgenesis, which was carried out by BestGene (http://www.thebestgene.com/). Transgenes were integrated into the genomic cytological position 53B2.

#### Generation of *flam* knockout line by CRISPR/Cas9

To remove the entire promoter region of *flamenco* we decided to delete the specific locus ChrX:21631436-21632105 (-455; +215 around TSS). To do so, we used two flanking sgRNAs. The two gRNAs were designed using CRISPR optimal target finder^82^. The individual guide sequences are listed in the **Table S3**. Bolded sequences correspond to each sgRNA. Those primers were used on a pCFD5 template. The resulting PCR product was assembled with the linearised pCFD5 backbone in a single Gibson Assembly reaction. Then the plasmid was injected in *vas*-Cas9 embryos (BestGene, BL#51324). To establish KO lines, molecular characterisation of target loci was performed as described^83^. Briefly, genomic DNA was extracted from individual larvae using QuickExtract DNA Extraction Solution (Cambio) according to the supplier’s instructions. Mutant flies were selected by PCR reactions. Exact deletion of *flam* was precisely identified by PCR and sanger sequencing once homozygote mutant fly lines were obtained (primers listed in **Table S3**).

#### Mitotic clones of *tj* mutant or knockdown analysis

For mitotic clones of tj mutant, the *Drosophila* strain *hsFlp; FRT40A - nls-RFP*; was crossed with the *tjeo2 FRT40A/CyO, bw* strain. For mitotic clones of *tj* knockdown, *y,w,HS:flp122/+;Act:FRTstopFRTGal4, UAS:GFP/CyO;* flies were crossed with lines carrying shRNAs against *tj* or *w* . All these flies were maintained at 25°C on a rich medium. Three heat shocks at 37°C for 1 h were applied to the F1 progeny, one day before eclosion (pupal stage), on the day of eclosion and one day after. The following day, ovaries were dissected and fixed for either FISH or IF experiments.

#### β-Gal staining on ovarioles

Ovaries from 5-day-old flies were dissected in PBS, kept on ice, fixed in 0.2% glutaraldehyde/2% formaldehyde/PBS at room temperature for 5 min and rinsed three times with PBS. They were then incubated in staining solution (1x PBS pH7.5, 1mM MgCl2, 4mM potassium ferricyanide, 4mM potassium ferrocyanide, 1% Triton, 2.7mg/ml X-Gal) at 37°C for 1h^69^.

#### FISH and immunostaining

Five to ten ovary pairs were dissected into ice-cold PBS and fixed in formaldehyde solution (4% formaldehyde, 0.3% TritonX-100 in PBS) for 20 min at RT with agitation. The fixed ovaries were then washed three times for 10 min in 0.3% Triton X-100/PBS and permeabilized overnight at 4°C in 70% ethanol. For probe hybridization, permeabilized ovaries were first rehydrated for 5 min in RNA FISH wash buffer (10% (v/w) formamide in 2x SSC). The ovaries were then resuspended in 50 μl hybridization buffer (10% (v/w) dextran sulphate and 10% (v/w) formamide in 2x SSC). Subsequently, 1.5 μL of a homemade *flam* RNA probe labelled with Atto-488 (Sigma, ref 41698) or Atto-550 (Lumiprobe, ref 92835) was added with or without primary antibody. The samples were incubated at 37°C with agitation. The ovaries were then rinsed twice for 15 min at RT in RNA FISH wash buffer. For FISH-IF, incubation with a secondary antibody conjugated to either Alexa-488, Cy3, or Cy5, was performed in 2x SSC for 1 h 30 at room temperature. After two washes in 2x SSC, DNA was stained with DAPI (1/500). The tissues were then mounted between a slide and coverslip using VectaShield mounting medium (Vector laboratories).

#### Immunofluorescence

Ovaries from 3- to 5-day-old flies were dissected in Schneider’s *Drosophila* Medium, fixed in 4% formaldehyde/PBS for 20 min, rinsed three times with PBT (× 1 PBS, 0.1% Triton, 1% BSA), and incubated in PBT for at least 1 h 30. They were then incubated with primary antibodies overnight. After 3 washes in PBT, ovaries were incubated with the corresponding secondary antibodies (1:1,000), conjugated to Alexa-488, Cy3, or Cy5, respectively, for 90 min. After two washes, DNA was stained with DAPI. The tissues were then mounted between a slide and coverslip using VectaShield mounting medium (Vector laboratories). All antibodies used for immunostaining (IF) along with probe sequences for in situ hybridization (FISH) are listed in **Table S3**. FISH and IF experiments were imaged using Zeiss LSM 800 or LSM 980 Airyscan confocal with a 20x and 40x objectives and analyzed using OMERO figure software.

#### TRAP experiment and RNA sequencing analysis

The TRAP experiment was described by Vachias and colleagues ^70^ and data was publicly available (see **Table S4**). Quality control of the sequencing data was evaluated using FastQC software. High quality reads were mapped to the *Drosophila melanogaster* dm6 reference genome using bowtie2 with default parameters^86^. Reads per gene were counted using HTSeqcount^87^. Normalization and differential gene expression analysis were performed using DESeq2^73^. Only genes with an adjusted p-value <0.05 were considered as differentially expressed between the two conditions. Gene expression data from Tj positive cells were compared to Nanos dataset by applying a fold change ≥ 2, *p* < 0.05 in order to generate list of enriched somatic genes. We computed and compared GO biological processes and molecular functions from this list using an R package cluster profiler from BioConductor^88^.

#### Small RNA sequencing

Total RNA was extracted from 80–100 pairs of ovaries from 3- to 5-day-old flies (for the analysis of piRNA production in somatic follicle cells) using TRIzol reagent (Invitrogen). After 2S rRNA depletion, deep sequencing of 18–30-nt small RNAs was performed by Fasteris S.A. (Geneva/CH) (**Table S4**).

#### Small RNA-seq analysis

Illumina small RNA-Seq reads were analysed with the small RNA pipeline sRNAPipe^84^ to map them to various genomic sequence categories of the *D. melanogaster* genome (release 6.03). All libraries were normalized per million of either genome-mapped reads or unique genome-mapping piRNAs (0 mismatches). All the analyses were performed using genome-unique piRNAs mapped to piRNA clusters, as defined by Brennecke and colleagues ^6^ (no mismatch allowed). The size profile for *flam* piRNA cluster was obtained by extracting the reads of 23–29 nt sequences uniquely mapped to this locus. Subsequent quantification of reads mapping to 1-kb tiles was done using bedtools, while relative quantification and plotting were done in R. Briefly, small RNA-seq tile signal was normalized to the estimated mappability scores for each 1-kb window. A pseudo-count of 1 was then added to each tile value and log2(fold-change) values relative to control samples was performed.

#### ATAC-seq data analysis

The read quality was assessed with FastQC (v0.11.8). The paired-end reads were trimmed of adapter sequences TCGTCGGCAGCGTCAGATGTGTATAAGAGACAG and GTCTCGTGGGCTCGGAGATGTGTATAAGAGACAG using Cutadapt tool (v1.18; default parameters)^64^. Burrows-Wheeler Aligner (BWA) (v0.7.17, bwa mem -M -t 4)^65^ was used to align the trimmed paired reads to 2014 (BDGP Release 6 + ISO1 MT/dm6) assembly of the *D. melanogaster* genome. Duplicates were marked using Picard tool (v2.9.0) (MarkDuplicates, validation stringency=lenient). SAMtools (v1.9) was used for indexing and filtering^65^. The quality metrics for the aligned ATAC-seq reads were assessed using ataqv (v1.0.0) (https://github.com/ParkerLab/ataqv)^66^. The ATAC-seq peaks were called with MACS2 (v2.1.1.20160309) using --nomodel --shift -37 --extsize 73 parameters and FDR cut-off q ≤ 0.05^67^. RPKM normalized bigWig files were generated using deepTools (v3.5.1, bamCoverage)^68^. Data were visualized using the UCSC Genome Browser^69^.

#### RNA-seq data analysis

The reads were trimmed of adapter sequences using Cutadapt tool (v1.18; -m 1 specified to not have reads trimmed to zero)^64^. The trimmed reads were aligned to the genome assemblies using the RNA-seq aligner STAR (v2.7.3a)^70^. Gene counts were calculated with the featureCounts tool (Subread package v1.5.3; -s 2 for stranded libraries prepared by dUTP method)^71^. *D. melanogaster* gene annotations were taken from Drosophila_melanogaster.BDGP6.28.100.gtf in the Ensembl genome database^72^ and annotations of orthologous *D. yakuba* genes were derived from *D. melanogaster* proteins mapped onto *D. yakuba* genome by chained tBLASTn (blastDm2FB in UCSC Table Browser). The read depth-normalized ovary RNA-seq counts were then normalized to species gene lengths. Consensus *Drosophila* transposon sequences were taken from (https://github.com/bergmanlab/transposons). Samtools (v1.9) was used to merge bam files from replicates^65^. Differential gene expression analysis was performed using DESeq2 (v1.30.1)^73^. The RPKM normalized bigWig files were generated for each strand using bamCoverage tool (--filterRNAstrand specified for dUTP stranded libraries) in deepTools (v3.5.1)^68^. Raw data for publicly available RNA-seq reads were downloaded from sources listed in **(Table S4)** and processed as above. Data were visualized using the UCSC Genome Browser^69^.

After quality control of reads using FASTQC (fastqcr_0.1.3), high quality reads were mapped to the *Drosophila melanogaster* dm6 reference genome (release 6_48 from flybase database), and consensus *Drosophila* transposon sequences (https://github.com/bergmanlab/transposons) using STAR (v2.7.8a) ^70^. Gene counts were calculated with the featureCounts tool (featureCounts v2.0.1). Normalization and differential gene expression analysis were performed using DESeq2^73^ (DESeq2_1.30.1). Volcano plots were generated using ggplot2 (v 3.3.3) in R (v 4.0.4).

#### ChIP-seq data analysis

The reads were trimmed of adapter sequences using Cutadapt tool (v1.18)^64^. The trimmed reads were aligned to the genome assemblies and TE consensus sequence assemblies (https://github.com/bergmanlab/drosophila-transposons/) using the Burrows-Wheeler Aligner (BWA) (v0.7.17, bwa aln)^74^. SAMtools (v1.9) was used for sorting, merging, and indexing^65^. ChIP-seq peaks were called using MACS3 (v3.0.0a6) callpeak command (-q 0.01)^67^ with data from ChIP input (the whole cell lysate) used as the control (-c). The reproducibility of ChIP-seq peaks between replicates was evaluated using the irreproducible discovery rate (IDR) tool (v2.0.2)^75^. The RPKM normalized bigWig files were generated using bamCoverage tool in deepTools (v3.5.1)^68^ with the parameter (-- extendReads 120) specified. The input-normalized bigWig files were generated using the bamCompare tool in the deepTools (v3.5.1)^68^. ChIP-seq heatmaps and profiles were generated using the plotHeatmap and plotProfile tools in the deepTools (v3.5.1)^68^. Raw data for publicly available ChIP-seq datasets were downloaded from sources listed in **(Table S4)** and processed as describe above. Data were visualized using the UCSC Genome Browser^69^.

Coverage plots over the TE consensus sequences were generated using the genomecov function in bedtools (v 2.26.0)^71^ with 1 nt bins and plotted with ggplot2 (v 3.5.1) in R (v 4.4.0). Normalised Tj ChIP coverage across the TE consensus sequences were created using the bamCompare function in deeptools (v 3.5.1) with RPKM normalisation and the log2FC operation. All TE consensus sequences were scaled to 5,000 bits using linear interpolation and the resulting coverage was plotted using ComplexHeatmap (v 2.20.0)^72^ in R (v 4.4.0).

#### Motif discovery and ATAC-seq peak conservation

*De novo* motif analysis of ChIP-seq peaks and known motif enrichments within peak centers were performed using the MEME-ChIP tool with the positional distribution of motifs calculated by CentriMo tool in MEME Suite (v5.4.1)^77^. Motif scanning was performed using FIMO tool (match p<0.001) in MEME Suite (v5.4.1) (https://meme-suite.org/meme/index.html)^76^. The list of *Drosophila* motifs was downloaded from the FlyFactorSurvey (http://pgfe.umassmed.edu/TFDBS/) database ^78^ and lists of known vertebrate motifs for peak central CentriMo enrichments were taken from JASPAR2022_CORE_vertebrates_non-redundant, EUKARYOTE/jolma2013, and MOUSE/uniprobe_mouse motif databases. fastaFromBed (v2.26.0) was used for conversion of bed file coordinates into fasta format using *D. melanogaster* BDGP6.28.dna.toplevel genome. Genomic loci were visualized on the UCSC genome browser^69^. Orthologous ATAC-seq peak regions between the species genomes were derived using the UCSC LiftOver tool (https://genome.ucsc.edu/cgi-bin/hgLiftOver)^79^. *D. melanogaster* ATAC-seq peaks were classified as conserved if the peak orthologous LiftOver regions in the *D. yakuba* genome overlapped with the *D. yakuba* ATAC-seq peaks.

### Data and code availability

RNA-seq and small-RNA-seq data generated in this study have been deposited to GEO database under series **GSE282040**.

#### Quantification and statistical analysis

Each experiment of qPCR data relative to *tomato* expression on fresh ovaries was performed 6 times with technical duplicates. Statistical differences were evaluated using Kruskal-Wallis non-parametric test, with corrections for multiple comparisons (Holm– Bonferroni adjustment method). Each experiment of qPCR relative to TE expression on ovaries was performed 3 times with technical duplicates. Fold changes of TE form *tj*-SKD compared to *w*-SKD were calculated. Statistical differences between *tj*-SKD and control (*w*-SKD) were evaluated using Wilcoxon non-parametric test using R software.

